# Low-dimensional Neural Codes Suppress Neuronal Noise and Extend the Working Memory Duration

**DOI:** 10.64898/2026.06.08.731010

**Authors:** Fatih Dinc, Chenzhang Feng, Marta Blanco-Pozo, Mathilde Papillon, Itamar Landau, Ruilin Shi, Mark J. Schnitzer, Peng Yuan, Nina Miolane

## Abstract

Neurons are noisy computational substrates, yet large neural populations achieve reliable computation. What determines the maximal duration that a noisy population can sustain working memory? We study this question with recurrent networks subject to stochastic noise and present three theoretical results. First, networks suppress independent neuronal noise when activity lies on a low-dimensional latent manifold. Second, this structure induces correlated noise across neurons, limiting the downstream information that can be extracted. Third, these effects yield an analytical bound on working memory duration that scales linearly with network size. We test these predictions using large-scale neocortical recordings, and provide a behavioral signature in mice consistent with the theory. Overall, noise suppression constitutes a key functional benefit of low-dimensional neural coding, with which large populations sustain reliable working memory over extended timescales.

## 1. Introduction

A computer holds in memory and processes gigabytes of data with perfect fidelity for hours. Yet a person asked to remember a long serial number will fail within seconds [1, 2, 3]. Similar limitations are observed across many species, from rodents to non-human primates [4, 5, 6, 7], inspiring over half a century of investigations [8, 9, 10]. Where does this limit originate from?

Prior work has proposed several cognitive and neural mechanisms underlying these limits, such as attentional capacity capped at a fixed number of slots [2, 11, 12], persistent recurrent activity limited by metabolic and resource constraints [6, 8, 13], activity-silent synaptic storage bounded by short-term plasticity timescales [14, 15], and brain oscillations constrained by intrinsic frequencies [16]. Each of these accounts, however, identifies a limit tied to a specific mechanism or implementation and thus may, in principle, be circumvented with a different strategy. In this work, we instead focus on a more fundamental and shared source of limitation, namely, the stochastic noise in neural activity.

This limitation rests on a fundamental difference between biological neural networks and computers. Computers are organized around a clear separation of memory, processing, and communication units [17], with errors regularly corrected to maintain signal fidelity, albeit at an increased energy cost [18]. The human brain, by contrast, computes in a distributed manner [19] with roughly 80 billion densely packed neurons [20]. Synaptic connections are continually remodeled [21], and neural activity is subject to constant stochastic fluctuations, *e*.*g*., those arising from intrinsic biophysical processes [22, 23] and the barrage of noisy signals from thousands of presynaptic inputs [24, 25]. As a result, alongside the signal that neurons transmit, they inevitably also transmit noise, producing irreducible trial-to-trial variability in their activities [26, 27]. In this work, we show that the resulting stochastic noise imposes a quantifiable limit on the duration of reliable computation involving working memory.

### Contributions

We introduce an analytical framework for noise propagation in a broad class of biologically plausible recurrent neural networks and establish three theoretical results (Fig. 1): i) Commonly studied low-dimensional codes suppress noise injected into neural activities, whereas high-dimensional codes do not (Proposition 1 and Interpretation 1). ii) After propagating through the same synaptic weights over time, independent noise becomes correlated trial-to-trial variability in neural activities. In turn, any useful downstream decoder picks up this correlated noise, limiting extractable information (Propositions 2, 3 and Corollary 1). iii) Latent noise accumulates and bounds reliable working memory computations to timescales that scale linearly with the network size (Proposition 4 and Corollary 2). We test these predictions on task-trained artificial networks and large-scale neocortical recordings [28]. In the end, we conclude with a novel, falsifiable hypothesis about how biological networks may code in two distinct regimes and provide preliminary behavioral signature consistent with this interpretation from mice trained on working memory tasks.

**Figure 1.**
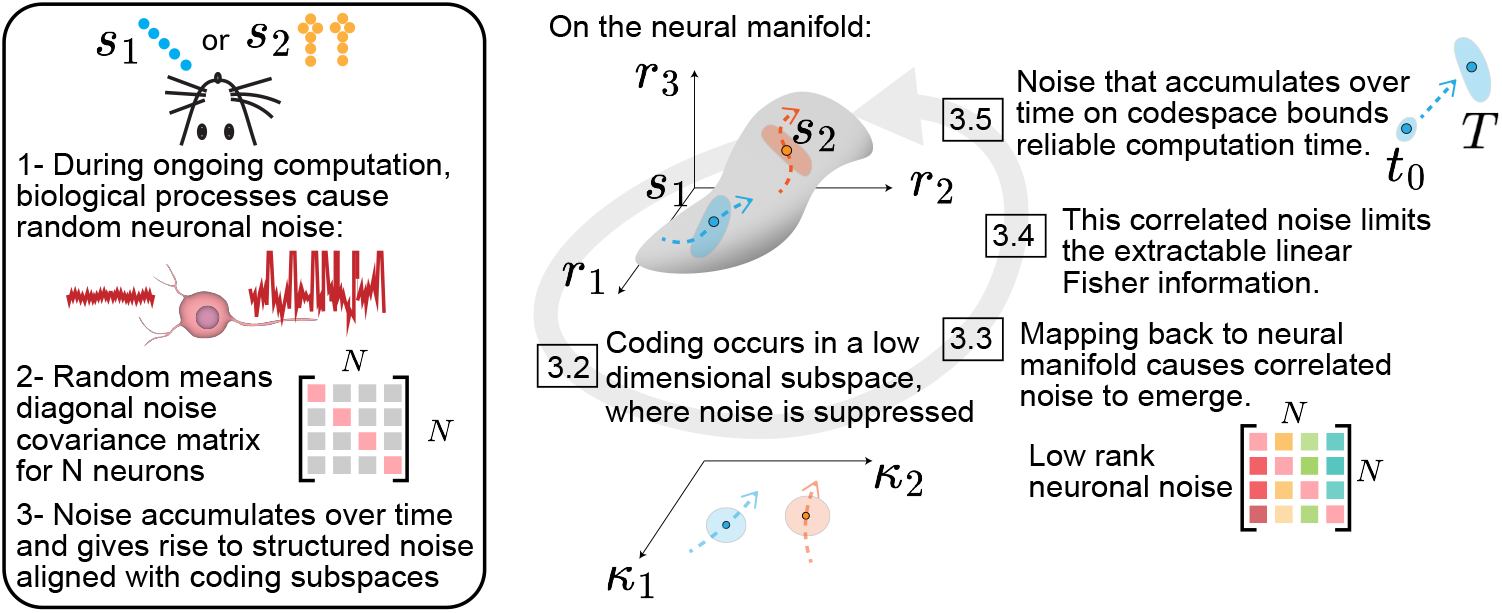
Summary of our theoretical results. *Left*. Neuronal noise is initially independent across neurons (1), yielding a diagonal *N × N* covariance matrix (2). Over time, it accumulates through the same dynamical system as the signal and aligns with the coding subspace (3). *Right*. On the neural manifold, computation is confined to a low-dimensional subspace spanned by *κ*_1_, *κ*_2_, where independent noise is suppressed (3.2). Mapping latent noise back onto the manifold, however, generates low-rank correlated noise (3.3) (pictured as overlapping clouds around two cues *s*_1_ and *s*_2_), matching the signal subspace. The correlated noise cannot be eliminated by pooling more neurons, limiting extractable information (3.4). As this correlated noise accumulates in the latent variables over time, the encoded state becomes increasingly uncertain, bounding reliable computation to a finite window *T* (3.5).

## 2 Related Works

### Variability in synaptic parameters

Synaptic weights undergo continual changes due to plasticity and turnover, altering the tuning properties of individual neurons, a phenomenon broadly termed representational drift [29]. Drift has been observed across a wide range of brain regions and species [30, 31, 32]. Several theoretical accounts propose that such drift reflects ongoing circuit plasticity and may contribute to continual learning by allowing neural circuits to remain malleable [33]. Empirically, some regions are better at maintaining stable population-level codes than others [28, 34, 35]. Understanding when and why this robustness emerges has long been studied [36, 37, 38].

### Noise in neural dynamics

A second, distinct source of noise arises from the randomness of neuronal processes themselves: independent noise injected into every neuron at every moment [22, 23, 24, 25]. Prior work has established that when such noise is correlated across neural populations, the information extractable by linear decoders saturates to a finite value even for infinitely large networks [26, 27, 39, 40, 41]. Large-scale recordings have confirmed the presence of such information-limiting correlations in neocortex [41, 42, 43]. How encoding fidelity evolves over time under constant noise injection, however, remains understudied. State-of-the-art analytical results so far are limited to linear networks [44], where the dynamics reduce to a well-known Ornstein-Uhlenbeck process [45]. Extending these calculations to nonlinear networks, where the noise covariance does not admit a closed-form Lyapunov solution, is one primary technical contribution of the present work.

### Low-dimensional neural code

Neural computation can often be described by a small number of latent variables [46, 47, 48] (though some cases call for high-dimensional codes [49, 50, 51, 52, 53]). This observation has been confirmed across species, brain regions, and behavioral paradigms, *i*.*e*., from motor cortex during reaching [54] to prefrontal cortex during decision-making [55], and from attractor dynamics underlying memory [56, 57] to flexible multi-task computation [58]. Theoretically, low-rank recurrent networks have provided mechanistic accounts of this low-dimensional activity [59, 60, 61, 62, 63] and offer a principled methodology for studying neural computation.

### Limits of working memory

The working memory capacity is differentially limited across species. In humans, short-term retention decays within seconds and is restricted to a few items [1, 2, 3, 11, 12]. Similar behavioral limits are evident in rodents [7], non-human primates [64], dolphins [65], pigeons [66], and honeybees [67]. Potential mechanisms underlying these limits have been studied extensively [68]. Recently, taking a dynamical systems perspective and assuming a low-dimensional neural code, Dinc et al. [62] showed that ongoing plasticity in synapses limits reliable computation with working memory to timescales scaling as 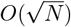, where *N* is the number of neurons in the network. Neuronal noise accumulation sets a secondary limit on information fidelity, which has not been understood yet and constitutes the focus of the present work.

## 3 Theory

### 3.1 Mathematical and conceptual setup

We study a biologically plausible class of nonlinear RNNs that unifies commonly studied architectures in systems neuroscience, including low-rank RNNs and state space models (see Appendix S1):

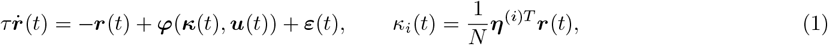

where *τ* ∈ ℝ^+^ is the neuronal timescale (milliseconds), ***r***(*t*) ∈ ℝ^*N*^ are population firing rates, ***κ***(*t*) ∈ ℝ^*K*^ are *K*-dimensional latent variables with linearly independent encoding vectors ***η***^(*i*)^ ∈ ℝ^*N*^ for 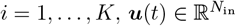 are external inputs, and ***ε***(*t*) ∈ ℝ^*N*^ is noise injected independent and identically distributed (iid) into each neuron at every moment. Without loss of generality, we assume ***η***^(*i*)*T*^ ***η***^(*j*)^ = *Nδ*_*ij*_ and refer to this as the canonical basis (proven in Appendix S2.1). This definition extends the noiseless construction of [62] by introducing ***ε***(*t*). With that, noise in neural activity immediately propagates into the latent variables:

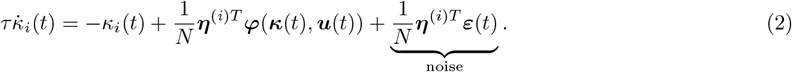

To enable analytical progress in nonlinear RNNs, we introduce one further ingredient. A key empirical fact about biological brains is that behavioral and neural timescales are separated by three orders of magnitude [69]: neurons fire on millisecond timescales while behavior and working memory unfold over seconds. We elevate this observation to a formal assumption:

#### Definition 1

(Separation of timescales). Latent dynamics evolve on an effective timescale:

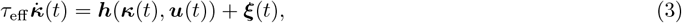

where 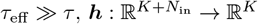 is the deterministic latent vector field, and ***ξ***(*t*) is the induced latent noise. The ratio *β* = *τ*_eff_ */τ* ≫ 1 is a large order parameter.

Realistically, *β* ~ 10^3^ to turn *O*(*ms*) neuronal time constant *τ* into *O*(*s*) behavioral timescale *τ*_eff_. This assumption will allow latent variables to evolve nonlinearly while we study small linear fluctuations around these mean trajectories.

#### Overview of theoretical results

Our central insight is a duality governing noise and signal. The same low-dimensional coding subspace that suppresses independent neuronal noise also forces that noise to re-emerge as correlated fluctuations aligned with the coding subspace. We formalize this in four propositions, illustrated in Fig. 1, culminating in a fundamental and quantitative bound on the duration of reliable computation under constant noise injection.

### 3.2 Low-dimensionality of latent variables enables noise suppression

We start by formalizing the noise terms with Wiener processes [45]:

##### Proposition 1

(Noise suppression in latent variables). *Under the assumption that injected noise is iid across neurons with zero mean, such that cumulative injected noise is Gaussian with variance* 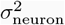 *per unit time, we arrive at stochastic differential equations (SDEs) [45]:*

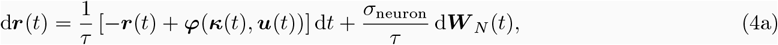

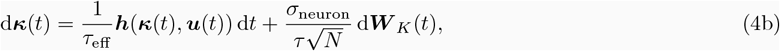

*where* d***W*** _*N*_ ∈ ℝ^*N*^ *and* d***W*** _*K*_ ∈ ℝ^*K*^ *are standard Wiener processes [45]*.

Note that ***W*** _*K*_ is the projection of ***W*** _*N*_ onto the latent subspace, coupling the noise dynamics between ***r*** and ***κ***. The iid Gaussian assumption, also assumed in earlier work [44], can be interpreted as coarse-graining many small stochastic contributions from channel noise, synaptic fluctuations, and presynaptic input variability into an effective Wiener process. The proof is presented in Appendix S2.1 and follows directly from Eq. (1) by applying the iid and Gaussianity assumptions and using the canonical basis property ***η***^(*i*)*T*^ ***η***^(*j*)^ = *Nδ*_*ij*_.

The key insight of Proposition 1 is that noise injected into neurons propagates into latent variables with an *O*(*N*^−0.5^) suppression, as shown in Eq. (4b). This is a direct consequence of the latent variable definition in Eq. (1), but this definition is self-consistent only if *K* ≪ *N*. When *K* ~ *N*, which is referred to as the extensive regime that can become chaotic [51, 70], latent variables in Eq. (1) need to be updated with 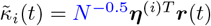, eliminating the 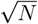 noise suppression of Proposition 1 entirely (see Appendix S2.2 for the derivations). This tradeoff reveals a key advantage of low-dimensional latent codes:

##### Interpretation 1

(Noise suppression with low-dimensional latent variables). *Biological networks may exploit low-dimensional latent variables precisely because population size then suppresses random noise by a factor of* 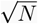. *This in turn enables reliable computation over the long timescales characteristic of working memory. In contrast, this suppression is absent in high-dimensional coding regimes, where an increase in population size cannot compensate for the injected noise*.

### 3.3 Correlated noise emerges in a low-rank subspace

Our goal is to characterize the noise structure in neural activity at a given time *t*. Solving Eq. (4a) yields three contributions to *r*_*j*_(*t*):

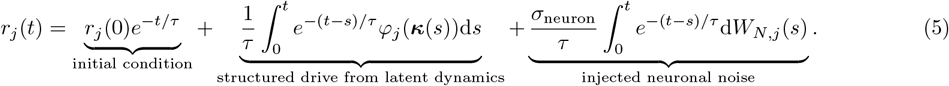

The first term is deterministic and vanishes on the fast timescale *τ*. The second captures the structured drive from latent dynamics, inheriting stochasticity from Eq. (4b). The third reflects independently injected neuronal noise. Because the second and third terms share the same underlying noise sources, they can interact and produce high-dimensional covariance structure. Characterizing this in full generality requires assumptions on ***φ***. Prior work obtained exact results by restricting to linear latent dynamics [44], where the problem reduces to the well-known Ornstein-Uhlenbeck process [45]. The following proposition makes progress by utilizing the timescale separation in Definition 1:

##### Proposition 2

(Diagonal plus low-rank noise covariance). *Assume that (i) φ*_*i*_ *is bounded and Lipschitz continuous for i* = 1, …, *N, (ii) β* = *τ*_eff_ */τ* ≫ 1 *with N* → ∞, *and (iii) small latent fluctuations (Proposition 1) permit approximating φ*(***κ***(*t*)) *via a first-order Taylor expansion around* ***φ***(𝔼[***κ***(*t*)]). *Then, for t* ~ *O*(*τ*_eff_), *the noise covariance of neural activities becomes:*

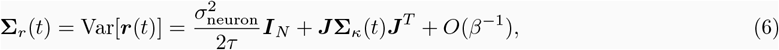

*where* ***J*** ∈ ℝ^*N ×K*^ *has entries J*_*ji*_ = ∂*φ*_*j*_*/*∂*κ*_*i*_|_𝔼[***κ***(*t*)]_ *and* **Σ**_*κ*_(*t*) = Var[***κ***(*t*)]. *The first term is diagonal (independent noise), the second is at most rank K (correlated noise)*.

The proof is given in Appendix S2.3. The key step exploits the timescale separation, *i*.*e*., since ***κ***(*s*) evolves on the slow timescale *τ*_eff_ ≫ *τ*, it is approximately frozen over the window *O*(*τ*) on which *e*^−(*t*−*s*)*/τ*^ is nonzero. This approximation decouples the structured latent drive from the injected noise to *O*(*β*^−1^) and, after a linearization of ***φ*** around 𝔼[***κ***(*t*)], yields Eq. (6).

Crucially, this diagonal plus low-rank structure arises without any assumption on the latent dynamics. Furthermore, while the noise increments d***W*** are Gaussian by construction, the accumulated noise in ***r***(*t*) need not be due to nonlinear ***φ***. Eq. (6) therefore suggests a model-free method for estimating latent dimensionality: fit a low-rank plus diagonal matrix to the empirical noise covariance without assuming any prior distribution on the accumulated noise. This is distinct from standard factor analysis, which assumes a generative model (*i*.*e*., Gaussian [71]) of the structured noise. We apply this method to task-trained RNNs and neocortical recordings in Section 4.2.

### 3.4 Noise in latent variables limits linear Fisher information

To study the coding fidelity despite noise, we consider a canonical binary cue discrimination task. On each trial, the network receives one of two stimuli *s*_1_ or *s*_2_ (cf. Fig. 1), and population activity ***r***(*t*) fluctuates around a cue-dependent mean 𝔼[***r***|*s*_1*/*2_] with trial-to-trial variability described by Proposition 2. A downstream observer decodes which cue was presented from the activity of *N*_obs_ ≪ *N* neurons via a linear decoder *ŝ*(*t*) = ***w***^*T*^ ***r***(*t*). To make all subsequent scaling arguments well-defined (see Appendix S2.4), the weights are normalized to 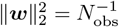. Defining *δ****r*** = *E*[***r***|*s*_2_] − *E*[***r***|*s*_1_], we can quantify the (local) coding fidelity with the following proposition:

##### Proposition 3

(Information-limiting latent noise). *Consider a linear decoder and assume a small separation between cues s*_1_ − *s*_2_ = *δs such that Eq*. (6) *has roughly the same parameter values across both stimuli, with* ***J*** *replaced with its counterpart* ***J*** _obs_ *defined over observed neurons. Then, the following statements are true:*

1. *The variance of the estimated cue ŝ*(*t*) *around s*_1*/*2_ *decomposes as:*

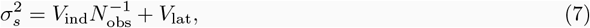 *where V*_ind_ *>* 0 *reflects independently injected neuronal noise and V*_lat_ ≥ 0 *captures the accumulated latent noise. By construction, V*_ind_ *and V*_lat_ *do not scale with N*_obs_.
2. *The linear Fisher information follows the well-known saturation curve [39, 42]:*

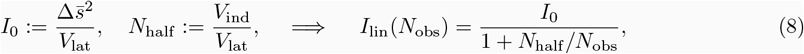 *where* 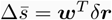 *is the separation of the decoded signal in neural activities*.

The key quantity in this proposition is 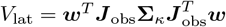, which represents the accumulated latent noise 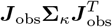 from Eq. (6)) projected onto the decoder weights ***w***. Then, for *V*_lat ≠_ 0 in Eq. (8), Fisher information saturates to a finite value *I*_0_. As *V*_lat_ → 0, both *I*_0_ and *N*_half_ diverge, recovering the classical result that information grows without bound in the absence of correlated noise aligned with the coding direction [39]. The following corollary clarifies that any decoder that locally separates the signal (*i*.*e*., 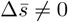) is subject to *V*_lat_ *>* 0:

##### Corollary 1

(Coding dimensions overlap with the noise subspace). *Any decoder with* 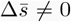 *must satisfy* 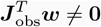. *Since* ***J*** _obs_ *also generates the correlated noise via Eq*. (6), *any such linear decoder, including the optimal one, incurs V*_lat_ *>* 0 *and the Fisher information saturates following Eq*. (8) *as N*_obs_ → ∞.

The proofs of Proposition 3 and Corollary 1 are given in Appendix S2.4. Importantly, these results are inherently local, as the noise covariance in Eq. (6) follows from a first-order Taylor expansion of ***φ*** around 𝔼[***κ***(*t*)]. Consequently, Corollary 1 requires *δ****r*** ∈ col(***J*** _obs_), which holds for *δs* → 0 but not globally. For commonly studied low-rank RNNs [59, 61] with globally linear coding manifolds, this condition holds everywhere, and information is therefore always limited. In contrast, with curved manifolds, *δ****r*** can remain orthogonal to col(***J*** _obs_), which could in principle allow *V*_lat_ = 0 for a binary decoder. In words, a key prediction arising from Proposition 3 is that curvature in neural manifolds can mitigate information limitation in downstream decoders.

### 3.5 Latent origins of accumulated noise

Even in simple binary decision making, most neurons have activities correlated with task-related variables [72]. Yet, if extractable information is severely limited and only a fraction of neurons is sufficient to communicate network computations, why would thousands of cortical neurons be recruited during simple tasks [28]? To answer this question, we now turn to the mechanistic origins of the latent noise **Σ**_*κ*_(*t*), which in turn incurs *V*_lat_ *>* 0:

##### Proposition 4

(Latent noise accumulation). *Consider a latent dynamical system as in Eq*. (4b) *and assume that* ***h*** *is smooth, with* ***h*** *itself and all its derivatives bounded by O*(1) *constants independently of N and β along any trajectory. Then, for t* ~ *O*(*τ*_eff_), *all eigenvalues of the latent noise covariance scale as:*

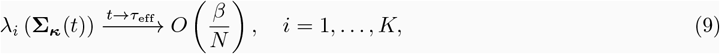

*where λ*_*i*_(**Σ**_***κ***_) *denotes the i-th eigenvalue of the latent noise covariance matrix* **Σ**_*κ*_.

The boundedness assumption on ***h*** and its derivatives is inspired by biological reality, namely that neurons fire in a physical world where firing rates change gradually and remain confined within physiological limits. The proof is given in Appendix S2.5 and follows a piecewise local linearization strategy of the latent dynamics. Combining Propositions 3 and 4, we arrive at a key corollary:

##### Corollary 2

(Duration of reliable computation). *Under the assumptions of Propositions 3 and 4, the reliable processing window scales as t* ∈ [0, *O*(*N*)], *and the number of neurons needed to extract half of the maximum information scales as N*_half_ ~ *O*(*N/β*).

This follows from plugging the *O*(*β/N*) scaling of Eq. (9) into Eq. (8), which gives *I*_0_ ~ *O*(*N/β*). A non-vanishing *I*_0_ then requires *β* ≪ *N*, establishing the *O*(*N*) bound on the processing window. For realistic numbers: *β* = 10^3^ converts millisecond neural timescales into second-scale behavioral ones, and *N* ~ 10^6^ (*e*.*g*., for mouse primary visual cortex [73]) gives *N*_half_ ~ 10^3^ consistent with empirical estimates [28, 42].

## 4 Experiments

### 4.1 After trial decay of Fisher information is consistent with seconds-long network timescales

A key assumption underlying all our results is the separation of timescales between neural and behavioral processes [69], namely *β* = *τ*_eff_ */τ* ≫ 1 in Eq. (3). Although biologically intuitive and broadly invoked, this assumption has, to our knowledge, never been directly tested in neural recordings. Existing justifications appeal to the general fact that neurons fire on *O*(*ms*) while behavior unfolds on *O*(*s*) [62, 69], but fall short of a direct test of this claim. Here, we propose and carry out a simple, parameter-free test. Eq. (3) implies that once the trial concludes, if ***h***(·) remains *O*(1) along the latent trajectory as assumed, any product of neural computation, *e*.*g*., the Fisher information of trial identity, must relax on the slow latent timescale *τ*_eff_ rather than the fast neuronal timescale *τ*. This prediction is directly testable, *i*.*e*., a post-trial decay constant of *O*(*s*) confirms *β* ~ 10^3^, whereas decay on calcium decay times of *<* 1*s* [74] would be a negative evidence.

We test this prediction by re-analyzing a previously published dataset with simultaneous recordings from up to eight neocortical regions in mice performing a Go/No-Go decision-making task [28]. In each trial, mice observe one of two stimuli for 2s, wait 0.5s, and then report by either licking a spout or abstaining. In line with prior work [28], we train linear decoders (Fig. S1) on neural activities at each time point to predict trial identity on correctly performed trials and estimate the associated Fisher information (details in Appendix S3). Across all eight regions, the Fisher information decays with *τ*_eff_ ≈ 2.5s (Figs. 2 and S2). This decay cannot be attributed to the autocorrelation times of individual neurons set by calcium decay, which remain sub-second for roughly 70% of the population (Fig. S3). Re-running the analysis on fast-decaying neurons only (*τ*_neur_ ≤ 0.5 s, where *τ*_neur_ denotes the estimated single-cell autocorrelation time; see Appendix S3) yields virtually the same results (Fig. S4). Finally, the licking and punishment probabilities decrease severely by the end of the trial [28], making the decay 10s after stimulus onset far less likely a byproduct of trivial trial correlates. Therefore, the extended decay is more likely a population-level effect consistent with *β* ≫ 1.

**Figure 2.**
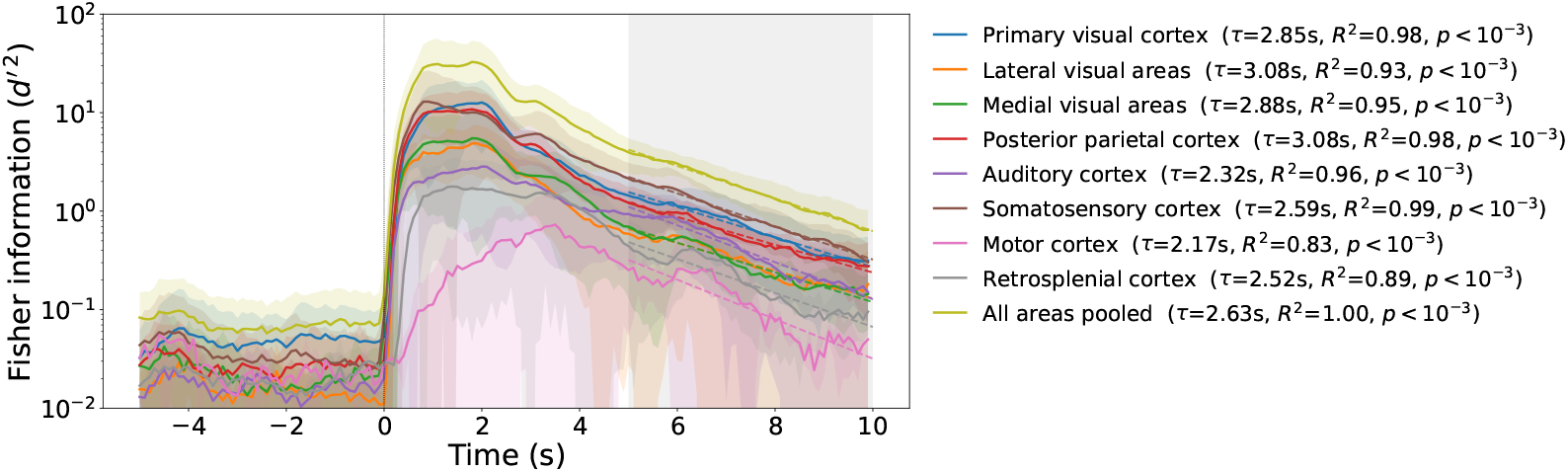
Fisher information of trial identity decays slowly after trial conclusion in the neocortex. Fisher information (*d*^*′* 2^) of trial identity is averaged across 100 random train/test splits per animal per time point. Solid traces show the cross-animal mean *d*^*′* 2^ for each cortical area, shaded bands denote *±*1 SD across animals. Time *t* = 0 s marks stimulus onset (vertical line); the shaded gray region (*t* = 5 s to *t* = 10 s) indicates the window after trial conclusion over which an exponential decay *d*^*′* 2^(*t*) ∝ exp(−*t/τ*) was fit to each area’s mean trace by linear regression on log *d*^*′* 2^. Dashed lines show the resulting fits; per-area decay constants *τ* are listed in the legend. Information decays on an effective timescale of *τ*_eff_ ≈ 2.5*s*.

### 4.2 Application of covariance matching to artificial networks and neocortical recordings

#### Learned low-rank computations in artificial neural networks

To validate covariance-matching process with simulations, we train vanilla RNNs and LSTMs on a canonical working memory task called *L*-bit flip-flop (for *L* ∈ *{*1, …, 5*}*, Appendix S4.2). RNNs often learn 2^*L*^ fixed point attractors to solve this task [75]. An effectively low-rank weight matrix would be expected to generate low-rank plus diagonal noise covariance around these attractors, with effective rank *K* ∈ [*L*, 2*L*] depending on the symmetry of the weight matrix [44]. To test it, we inject iid Gaussian noise at inference time, and fit a diagonal-plus-rank-*K* model to the resulting noise covariance matrices via alternating minimization (Appendix S4.1). We note that while low-rank RNNs are members of the network family in Eq. (1) by construction, full-rank RNNs and LSTMs are not. Yet, they may still effectively learn low-dimensional solutions that respect Eq. (6), which is what we test here.

For vanilla full-rank RNNs, the test correlation saturates sharply at rank *K* = *L* across all noise levels (Fig. S5, top row), consistent with the hypothesis that the latent dimensionality would be determined by the task requirements [52, 62]. For LSTMs, a low-dimensional correlated noise structure also emerges, though the saturation is less sharp and the estimated rank *K* can modestly exceed *L* at higher task complexity but remains *K* ≤ 2*L* (Fig. S5, bottom row). These results support the interpretation that the noise covariance carries a useful signal about latent dimensionality.

#### Cortical noise is consistent with tens to a hundred latent dimensions

We next apply this framework to calcium imaging recordings from mouse neocortex [28]. We estimated latent dimensionality across seven cortical regions using four complementary methods (Appendix S4): participation ratio on the full covariance matrix, participation ratio on the off-diagonal covariance matrix (diagonal set to zero, eigenvalues thresholded to be positive), covariance matching (CM), and factor analysis (FA).

The participation ratio on the full noise covariance is high across all areas, near the full recorded population size (Figs. 3 and S6). This is consistent with Eq. (6) if the diagonal independent-noise term dominates the full covariance, making the noise itself high-dimensional consistent with [49, 50]. Setting the diagonal to zero reduces the participation ratio substantially (Fig. 3), but the resulting estimate mixes the true low-rank correlated component with two noise sources due to finite-sample estimation and an artifact from removing the non-zero diagonal entries of the structured component.

**Figure 3.**
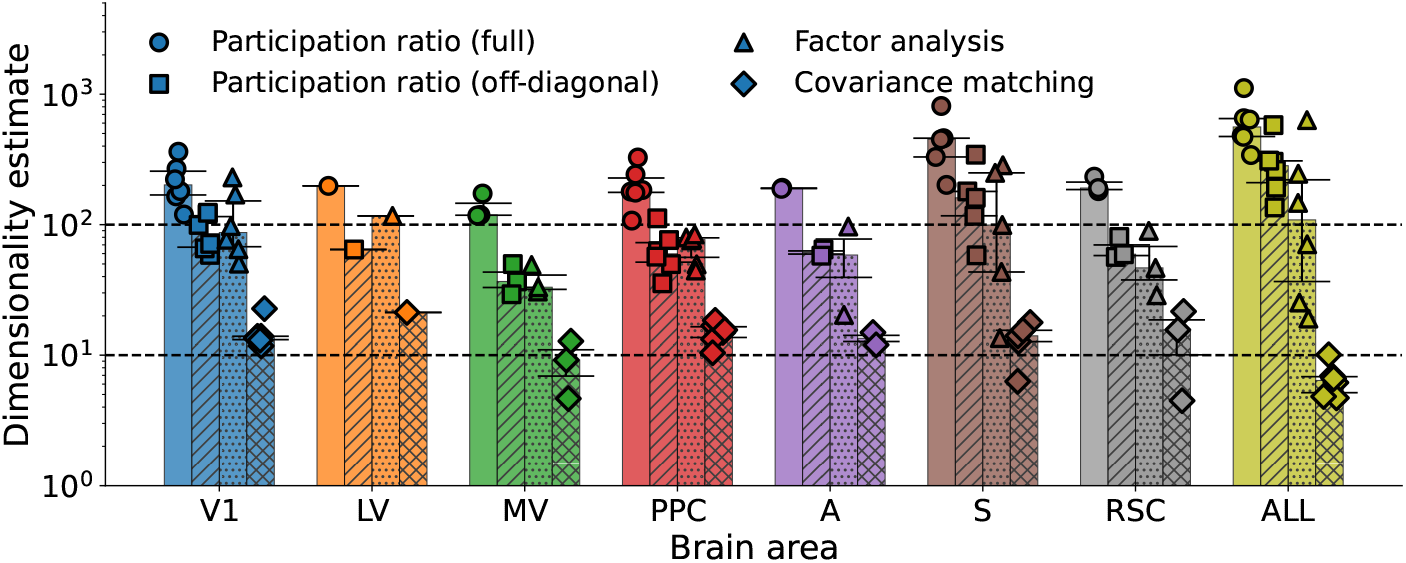
Cortical noise occupies tens to a hundred latent dimensions across areas. We estimated latent dimensionality across cortical areas using four methods: participation ratio computed on the full covariance matrix (full) vs when diagonal elements were set to zero and only positive eigenvalues used (off-diagonal), factor analysis (FA), and covariance matching (CM), details in Appendix S4. Bars show the median across animals (*n* = 6), error bars denote the interquartile range, and each dot represents an animal. The horizontal lines correspond to 10 and 100 latent dimensions. Brain regions are color-coded as in Fig. 2.

The covariance-matching estimates isolate the correlated component directly through cross-validated model selection. The test correlation saturates at rank *K* ∈ [10, 30], stable across the stimulus, delay, and response periods (Figs. 3 and S7). Notably, since covariance-matching identifies the dominant modes contributing to correlated noise, it can underestimate the true latent dimensionality (cf. Fig. 3, where dimensionality estimates using neurons pooled from all brain regions can become lower compared to those estimated within individual regions). We supplemented this method with a second method, *i*.*e*., factor analysis. FA yielded higher estimates in the range of tens to hundreds (Figs. 3 and S8), likely because the Gaussian generative model need not hold for noise accumulated through a nonlinear embedding. Overall, the correlated component of the noise covariance is consistent with a latent subspace *K* ∈ [10, 100]. This agrees with the findings of prior empirical works [28, 42, 76], in line with Interpretation 1 that noise suppression is effective when computation is confined to low-dimensional subspaces.

### 4.3 A behavioral test of memory retention in mice

Earlier work has predicted that in a network where synaptic connections randomly change, the parameters of the latent dynamical system exhibit 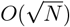 fluctuations, *i*.*e*., ***h*** in Eq. (3) would have 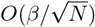 uncertainty, limiting reliable computation to behavioral timescales 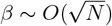. Combined with Proposition 4, this predicts the existence of two distinct coding regimes in biological neural networks (Fig. 4**A**). At short delays, working memory can be reliably maintained despite ongoing representational drift, whereas training at longer delays can selectively stabilize synaptic weights in task-relevant subpopulations. Once drift is suppressed, irreducible neuronal noise sets a second, harder limit scaling as *β* ~ *O*(*N*) (Corollary 2). We hypothesize that the transition between these regimes should manifest as a localized increase in learning difficulty. To test this, we trained mice on a two-alternative forced-choice delayed discrimination task with progressively increasing delay durations (Fig. S9). To reach the next level of delay, mice had to perform 70% correct in a session.

**Figure 4.**
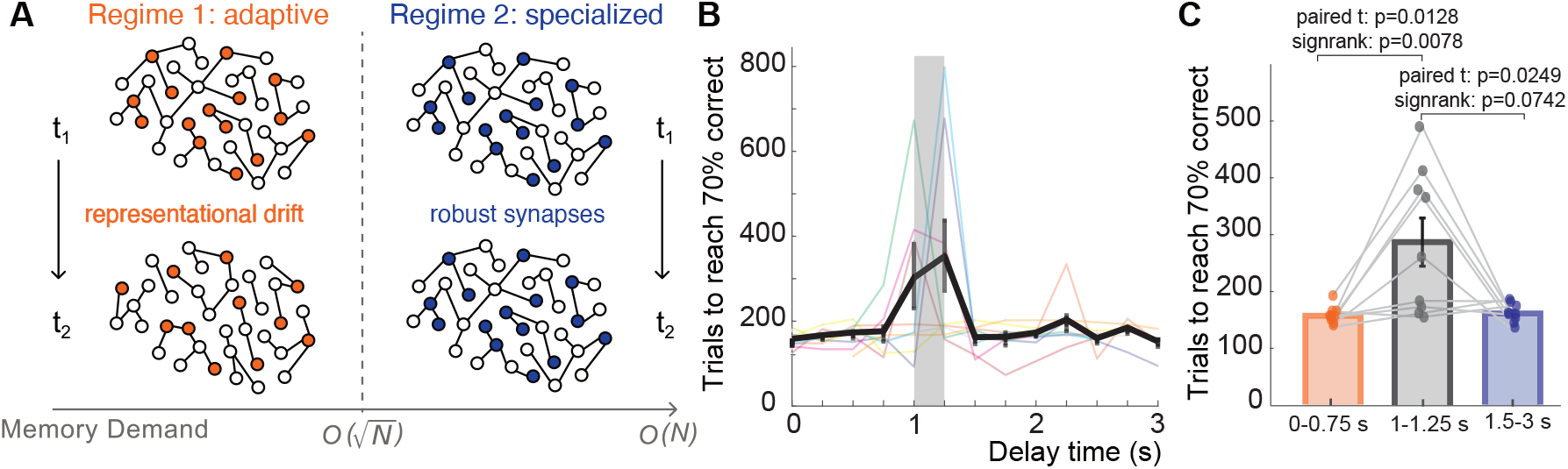
Mice trained on a working memory task reveal a behavioral signature of two distinct reliability regimes. **A** Schematic of the two theoretically predicted coding regimes. The transition between regimes predicts a localized increase in learning difficulty at the delay duration where the drift-imposed ceiling is reached. **B-C** To test this prediction, we trained mice on a two-alternative forced choice delayed discrimination task with the delay duration increasing in 0.25s increments over time (see Fig. S9 for the training paradigm). **B** Number of trials required for each mouse to reach 70% correct at each delay stage. Thin colored lines denote individual mice; the thick black line indicates the mean *±* s.e.m. The shaded region marks the 1-1.25 s delay interval. **C** Trials required to reach 70% correct for grouped delay values. Bars indicate mean *±* s.e.m.; dots represent individual mice; gray lines connect paired observations. n=9 mice.

As delays increased, mice required significantly more trials to reach criterion at the [1, 1.25]*s* delay intervals than at shorter delays (Fig. 4**B-C**, Cohen’s *d* = 1.06), consistent with a localized increase in learning difficulty at the predicted transition boundary. Critically, performance at delays beyond 1.25s recovered and required fewer trials to reach 70% correct than the transition interval, with values comparable to short delay (*p* = 0.508). We also noticed that a few mice showed no difficulty during the 1-1.25 s window. Interestingly, these mice tended to show superior learning efficiency during Stage 1 training (Fig. S10, though this mean difference did not reach significance, *p* = .152), suggesting that they may have already crossed the barrier in the initial learning of the task. As we will argue below, large-scale recordings can test these predictions at the level of individual variability. Overall, our results in Figs. 4, S9, and S10 were consistent with mice adopting a qualitatively different coding strategy in the noise-dominated regime.

Our framework predicts that once this strategy is adopted, generalization to longer delays should incur no further learning difficulties until the much larger *O*(*N*) bound is reached. In line with this hypothesis, once trained to stable performance at 3s delay, mice generalized to novel 6s delays without additional training (Fig. S11). Performance was preserved in a blocked setting (Fig. S11**A**, *p* = 0.563), where 6s delays were introduced at the end of a session. Similar results were obtained in the interleaved design (Fig. S11**B**, *p* = 0.313), where 6s delays were randomly introduced interleaved with the 3s delays. Together, these results provide convergent preliminary evidence that the localized learning difficulty reflects the predicted regime transition. We emphasize that behavioral results alone cannot prove the existence of the two coding regimes, as alternative explanations exist (cf. limitations below). They instead establish the two-regime hypothesis as a falsifiable prediction worth direct tests with large-scale neural recordings coupled with targeted behavioral experiments.

## 5 Discussion

In this work, we develop an analytical theory of noise propagation in a broad class of biologically plausible neural networks by studying linear perturbations around nonlinear latent trajectories. The key ingredient enabling this progress is the commonly invoked separation of timescales assumption (Definition 1), which we explicitly test in Fig. 2, to our knowledge for the first time. By establishing a second, softer bound on working memory duration alongside the drift-imposed bound of prior work [62], our framework predicts the existence of two distinct coding regimes in biological neural networks. We speculate that the transition between these regimes may hold the key to understanding how biological systems balance adaptive learning against stable working memory computations.

### Connections to systems neuroscience

Proposition 1 adds to existing theoretical accounts of low-dimensional population codes [48, 77, 78] by identifying noise suppression as a key functional benefit. Proposition 2 then predicts a measurable fingerprint. The noise covariance of neural activities takes a diagonal plus low-rank form whose rank equals the latent dimensionality, enabling dimensionality estimation from the noise rather than the signal (Fig. 3). This structure connects naturally to traditional accounts of information-limiting noise correlations, which focus on static sensory representations in feedforward networks [26, 27, 39, 79]. Proposition 3 extends this picture to recurrent networks without ongoing sensory drive, showing that the alignment between coding and noise subspaces arises naturally from latent variable structure. Finally, Proposition 4 shows that this latent noise accumulates over time and generates a fundamental bound on reliable working memory computation.

### Connections to attractor dynamics

The *O*(*N*) scaling of working memory duration is consistent with a rich literature on attractor networks, from the foundational work of Hopfield [80] to later studies of discrete and continuous attractors [13, 57, 81], where working memory capacity is known to scale linearly with network size in many specific cases. In this work, we derive this bound starting from first principles in a general latent variable framework, without assuming a particular attractor geometry or specialized network architecture, and establish the link to the latent dimensionality.

### Testable predictions

Our work yields three testable predictions. First, the machinery we used to prove Corollary 1 predicts that manifold curvature modulates information limitation. When the signal direction lies outside the tangent manifold, the correlations may become orthogonal to the signal. So, brain areas with curved coding geometries could exhibit weaker (or not at all) information saturation for categorical codes. Second, the low-rank plus diagonal covariance structure provides a model-agnostic tool for estimating latent dimensionality from noise. The covariance matching procedure in Fig. 3 is a first implementation. More principled methods would yield sharper estimates and a statistical test of the predicted structure.

Finally, the two coding regime hypothesis predicts that suppressing representational drift extends working memory timescales at the cost of representational flexibility, creating a fundamental tradeoff between stable retention and adaptive learning. Our behavioral results provide indirect evidence for this tradeoff, though large-scale neural datasets combined with targeted behaviors offer an immediate opportunity to test this hypothesis further. Moreover, direct causal tests via targeted perturbations of synaptic consolidation mechanisms are needed to establish it.

### Limitations and future work

Our work has several limitations. First, our theoretical results rest on a linearization of the latent dynamics around the mean trajectory (Lemma S2). This is valid when noise fluctuations are small, but may break down for networks operating far from this regime. A nonlinear treatment of large fluctuations remains an important open problem. Second, the timescale separation assumption (*β* ≫ 1) enables our analytical progress. We test this assumption empirically in Fig. 2, but it need not hold for all tasks and brain regions, *e*.*g*., fast sensorimotor tasks requiring decisions on millisecond timescales. Third, our experimental results rely on public calcium imaging datasets [28], which conflates neural timescales with indicator decay kinetics. Although our control analyses show that slowly decaying individual neurons do not explain the population-level decay (Fig. S4), replication with electrophysiology in more diverse datasets would provide a secondary test.

On the conceptual side, our framework applies most directly to low-dimensional coding regimes, where population averaging suppresses noise. A similar analysis for commonly studied high-dimensional codes suggested that random noise is not suppressed in latent variables. However, for general high-dimensional codes, neural populations may not revisit stereotypical activity states [82, 83], and this suppression mechanism may not even be needed, as the high-dimensional geometry itself may provide alternative means of separating signal from noise. Whether an analogous bound on working memory duration exists in such regimes, or whether high-dimensional codes circumvent the limits identified here through qualitatively different computational strategies, remains an open question.

Finally, the behavioral experiment is not a comprehensive characterization of the regime transition and cannot rule out alternative explanations. Other interpretations of the localized learning difficulty are possible, including a single-strategy capacity limit, task-restructuring effects, or transient motivational and attentional changes at the boundary delay. Distinguishing these requires neural recordings during the transition, which is beyond the scope of the present theoretical work. We deliberately frame Fig. 4 as a behavioral signature consistent with the theory, not as a definitive test, and view its primary value as motivating large-scale neural recordings during the predicted transition.

### Broader implications for artificial intelligence

Our framework connects to two lively debates in machine learning. The first one concerns in-weight vs in-context learning [84]. The two physical bounds derived here suggest a concrete asymmetry. The adaptive coding regime is analogous to the in-context learning, which uses existing strategies of the network adaptively. In contrast, test-time training that updates parameters [85] and creates task-specific structure is analogous to in-weight learning. While this analogy breaks at the substrate level, as test-time training changes parameters whereas in-context learning does not, the conceptual connections do exist. Studying how biological networks navigate this physical tradeoff may provide further insights into this debate. The second debate is about the use of physical computing systems. Analog, neuromorphic, and optical systems promise dramatic energy savings over digital architectures [18] but may reintroduce the very noise problems that digital abstraction was designed to eliminate. Biological neural networks have navigated this tradeoff for hundreds of millions of years, and the framework developed here characterizes one fundamental limit they face, offering a principled reference point for understanding how noisy physical substrates can sustain reliable computation.

## Acknowledgements

We thank Louis Van Langendonck and Geometric Intelligence lab members for their constructive inputs to a preliminary draft of this manuscript. This research was supported in part by grant NSF PHY-2309135 and the Gordon and Betty Moore Foundation Grant No. 2919.02 to the Kavli Institute for Theoretical Physics (KITP).

## S1 Extended mathematical background

### S1.1 Information-limiting noise correlations

Traditional work on how much information can be extracted from an observed population of neurons has primarily focused on static representations encoded in neural activities [26, 27, 39, 40, 79, 86]. To summarize these findings and motivate our extension to networks with recurrent dynamics, we consider a binary cue discrimination, which is probed with a standard set of tasks studied in systems neuroscience for more than three decades [87].

#### Fisher information associated with a binary cue discrimination task

We assume that activities from *N*_obs_ neurons are recorded while two distinct cues *s*_1,2_ are presented. Each presentation of a cue is a trial; within each trial, neural activities fluctuate around their mean, 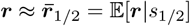, with trial-to-trial variability captured by the noise covariance matrices **Σ**_1*/*2_ = Var[***r***|*s*_1*/*2_]. In both cases, expectations are taken over trials. With two assumptions, namely that **Σ**_1*/*2_ = **Σ** is shared between classes and that the noise is Gaussian, the optimal readout becomes linear, ***w***^∗^ ∝ **Σ**^−1^**Δ*µ***, where 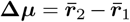 is the mean separation. Then, the (unitless) Fisher information extractable from the neural population takes the form [40]:

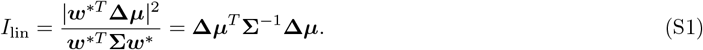

When the noise is uncorrelated 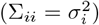, then 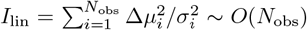 grows without bound as more neurons are observed, which simply means that pooling independent samples reduces error.

#### Information-limiting differential correlations

Earlier work has shown that this scaling is broken, and *I*_lin_ saturates to a finite limit, if and only if the noise covariance contains a component aligned with the signal direction [39]:

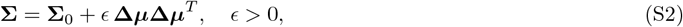

where **Σ**_0_ is the component that does not limit information. The rank-one term **Δ*µ*Δ*µ***^*T*^ is referred to as differential correlations: noise whose covariance is aligned with the signal direction **Δ*µ***, and which therefore cannot be averaged away by pooling more neurons. The existence of differential correlations has since been confirmed in large-scale neural recordings [41, 42, 43]. As a reflection of the low-dimensional structure in Eq. (S2), only a small number of dimensions of **Σ** have been relevant for cue discrimination in decoders trained on the activities of neurons across the full neocortex [28].

The origin of differential correlations, however, has remained poorly understood. Earlier work has shown that noise injected at the sensory periphery and propagated through a feedforward network naturally produces differential correlations [79]. This mechanism, however, cannot explain information limits in settings where no ongoing sensory drive is present, and the population must maintain a representation through recurrent dynamics alone. In this work, we extend this line of work by showing that differential correlations arise naturally in recurrently connected networks from accumulation of independent noise in low-dimensional latent variables.

### S1.2 A general model of computation in biological neural networks

We consider a broad class of biologically plausible neural networks:

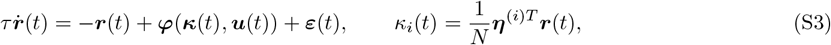

where ***r***(*t*) ∈ ℝ^*N*^ denotes the firing rates of *N* neurons, ***κ***(*t*) ∈ ℝ^*K*^ are *K*-dimensional latent variables, 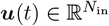 are external control inputs (*e*.*g*., task-dependent stimuli), and ***ε***(*t*) is independent and identical neuronal noise arising from biological processes. The scalar *τ >* 0 is the neuronal time constant, and ***η***^(*i*)^ ∈ ℝ^*N*^ is the encoding weight vector of the *i*-th latent variable, which linearly projects neural activity onto *κ*_*i*_. The embedding map 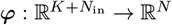 explains neural dynamics as driven by low-dimensional latent variables together with external inputs. Under standard assumptions on ***φ*** (*e*.*g*., non-polynomial nonlinearity [88, 89]), Eq. (1) becomes a universal approximator of dynamical systems without further constraints on the network architecture [62].

#### Population coding in *N* → ∞ limit

Focusing primarily on the noiseless setting (***ε***(*t*) = 0), Ref. [62] has shown that Eq. (1) recovers the commonly studied low-rank RNNs [59, 60, 61, 90] and state space models [91, 92, 93] as a special case while remaining agnostic to the specific network architecture across a broad, biologically plausible class of models. Without the need for further architectural specification, Eq. (1) induces the following latent dynamical system:

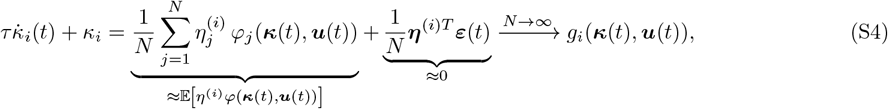

where 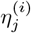 denotes the *j*-th entry of ***η***^(*i*)^, and *φ*_*j*_ is the *j*-th component of the embedding map, whereas (*η*^(*i*)^, *φ*) refer to the random variables from which these samples are drawn^1^. The *N* → ∞ limit is often referred to as the mean-field limit, commonly taken in low-rank RNN studies under the assumption that synaptic weights scale as *O*(*N*^−1^) [60] (though also see [51, 70] for a different, chaotic regime). In this limit, the behavior of the latent dynamical system is governed not by the individual neuronal parameters 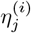 and *φ*_*j*_, but by the distribution 𝒫 over (*η*^(*i*)^, *φ*) from which they are drawn. The expectation is taken with respect to 𝒫 and defines the latent dynamics ***g***(***κ***(*t*), ***u***(*t*)). In this work, we use this mathematical limit whenever we refer to the population coding.

#### Two key sources of deviations in population dynamics

For finite *N*, there are two key sources of errors that describe the deviation of population dynamics from ***g***(***κ***(*t*), ***u***(*t*)), each associated with a term in Eq. (S4). The first term in Eq. (S4) describes the target behavior in the mean-field limit, whereas deviations arise from fluctuations in the synaptic parameters for finite *N*. This term is quenched, *i*.*e*., fixed once a network is actualized. Prior work has studied the effects arising from variations in this term assuming ***ε***(*t*) = 0 [62] and found that large populations of neurons are needed to accurately construct ***g***(***κ***(*t*), ***u***(*t*)) when latent variables evolve in longer timescales than *τ* (see next section). In this work, we study the stochastic second term, in which the random noise is injected at every time point to each individual neuron independently and regardless of the network architecture. Our goal is to characterize how this neuron-level noise propagates into the latent dynamics, accumulates over time, and ultimately limits the fidelity of computation over extended timescales.

### S1.3 Computation in extended timescales under noisy dynamics

To study computation over extended timescales, we assume that the latent dynamical system is trained to implement the following deterministic dynamics in the limit *N* → ∞:

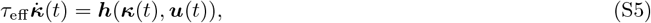

where *τ*_eff_ is the effective latent time scale underlying behavioral processes and ***h***(***κ***(*t*), ***u***(*t*)) describes the latent dynamics induced in the limit *N* → ∞. By assumption, ***h***(***κ***(*t*), ***u***(*t*)) does not depend on the ratio *β* = *τ*_eff_ */τ*. (As a side note, since *τ* ~ *O*(*ms*) and *τ*_eff_ ~ *O*(*s*), a realistic *β* ~ 10^3^ ≫ 1.) Rescaling time in Eq. (S5) as *t* → *t/τ*_eff_ yields scale-free dynamics, making *β* an order parameter that controls the timescale over which computation unfolds. Matching Eqs. (S4) and (S5), the target dynamics for finite *N* are approximated by:

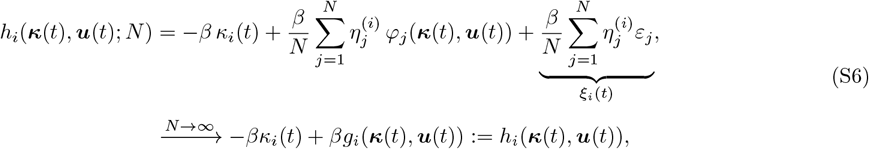

As discussed in earlier section, the finite-*N* deviations between ***h***(***κ***(*t*), ***u***(*t*); *N*) and ***h***(***κ***(*t*), ***u***(*t*)) arise from two distinct sources. For this work, we assume that synaptic weights are chosen more precisely and the finite-sample approximation error of 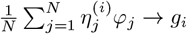 vanishes for finite *N*. Then, only injected noise *ξ*_*i*_(*t*) remains and cannot be eliminated by more precise synaptic weights, such that the latent dynamics for finite *N* follows:

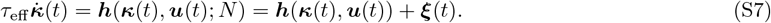

This equation constitutes the starting point for our subsequent analyses.

## S2 Theoretical details and proofs

### S2.1 GL(*K*) symmetry of encoding weights and proof of Proposition 1

In this section, we first introduce a simple Lemma and use it to prove Proposition 1.

#### Lemma S1

(Canonical normalization). *Without loss of generality, one can assume* ***η***^(*i*)*T*^ ***η***^(*j*)^ = *Nδ*_*ij*_ *as long as* ***η***^(*j*)^ *are linearly independent for j* = 1, …, *K*.

*Proof*. Let ***η*** ∈ ℝ^*K×N*^ denote the matrix whose *i*-th row is ***η***^(*i*)*T*^, so that the latent variables are 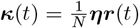. Under the transformation

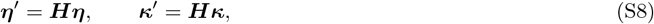

for any invertible ***H*** ∈ *GL*(*K*) (the group of *K × K* invertible matrices), the encoding definition is preserved since 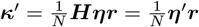. Then, the neural dynamics in Eq. (1) become:

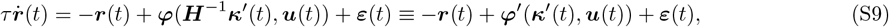

where ***φ***^*′*^(·, ***u***) = ***φ***(***H***^−1^·, ***u***) is a reparametrized embedding map. Since ***φ***^*′*^ is an equally valid map to ***φ***, the neural dynamics are invariant under this transformation. The Gram matrix ***G*** ∈ ℝ^*K×K*^ with entries *G*_*ij*_ = ***η***^(*i*)*T*^ ***η***^(*j*)^ transforms as ***G*** → ***HGH***^*T*^. Since the encoding vectors 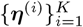 are assumed linearly independent, ***G*** is symmetric positive definite and ***G***^1*/*2^ exists. Choosing

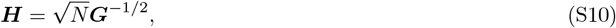

which is invertible since ***G*** is positive definite, and symmetric since ***G***^−1*/*2^ is symmetric, gives:

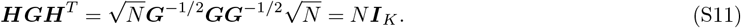

Hence the transformed encoding weights satisfy ***η***^*′*(*i*)*T*^ ***η***^*′*(*j*)^ = *Nδ*_*ij*_, completing the proof.

Now, for completeness, we restate Proposition 1 (with a small change in Eq. 4 that moves the timescales to the left hand side) before proving it:

*Proposition*. Under the assumption that injected noise is iid across neurons with zero mean, such that cumulative injected noise is Gaussian with variance 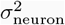 per unit time, we arrive at stochastic differential equations (SDEs) [45]:

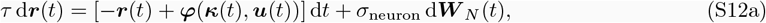

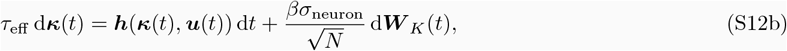

where d***W*** _*N*_ ∈ ℝ^*N*^ and d***W*** _*K*_ ∈ ℝ^*K*^ are standard Wiener processes [45].

*Proof*. Under the substitution of the Wiener Process, the latent noise term defined in Eq. (S6) satisfies:

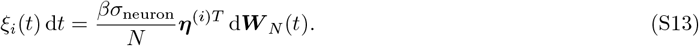

Under the canonical normalization, Eq. (S13) reduces to:

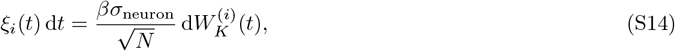

which follows from the fact that 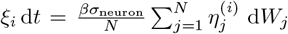 is a linear combination of independent Wiener increments, hence itself a Gaussian increment with variance:

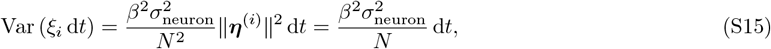

using ∥***η***^(*i*)^∥^2^ = *N* in the canonical basis. Similarly, for *i≠ k*:

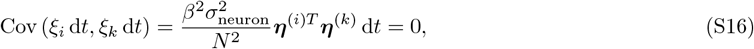

by orthogonality. This concludes the proof.

### S2.2 Neural noise in the extensive rank limit

To understand why the noise suppression is practically gone in the extensive limit *K* → *N*, we first consider a biologically motivated embedding map ***φ*** (noiseless and no inputs for simplicity) that acts on a vector of currents received by each neuron [62]. Specifically, neuron *j* receives *L* ~ *O*(1) distinct currents through distinct dendritic compartments, each written as a summation over latent variables:

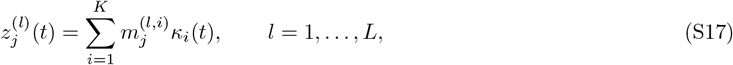

where ***m***^(*l,i*)^ ∈ ℝ^*N*^ are the embedding weights for the *l*-th current and *i*-th latent variable. The individual elements scale as 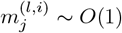, which results from the scaling of synaptic weights 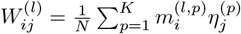. Specifically, the scaling 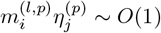 recovers the standard *O*(1*/N*) weight scaling of the mean-field limit *K* = *O*(1) [60] and the 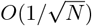 weight scaling of the extensive regime *K* = *O*(*N*) when latent variables are decoupled [51, 70]. The case *L* = 1 with a tanh(·) nonlinearity recovers the basic RNNs extensively studied in both limits [51, 60, 62, 70]. The embedding map then acts on the full current vector of neuron *j*:

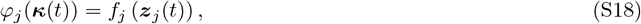

where *f*_*j*_ : ℝ^*L*^ → ℝ is an element-wise nonlinearity. For the network to operate without currents blowing up, each current must remain *O*(1). But, if we assume that finite groups of latent variables are responsible for independent parts of the computation, enforcing *κ*_*i*_ ~ *O*(1) leads to a contradiction when *K* → *N* for a generic construction. This is because, with 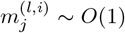, each current is a sum of *O*(*N*) iid *O*(1) terms, giving 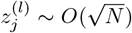. In the extensive limit *K* → *N*, this means that every current diverges, leading to blow-up in the network dynamics as *N* → ∞. One could technically still define latent variables using the saturation property of ***φ*** or by a specific code that constraints all *O*(*N* ^2^) interactions between latent variables to cancel each other out. However, the former leads to biologically implausible and non-physical infinite currents, whereas whether the latter can support useful computation remains unknown.

The commonly studied resolution comes from the observation that in the extensive limit, networks often enter a regime in which neural activities decorrelate across neurons [51, 70]:

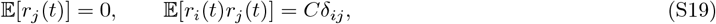

for some *C* ~ *O*(1). Since *r*_*j*_ are independent with zero mean and variance *C*, computing the latent variables under the 1*/N* normalization gives:

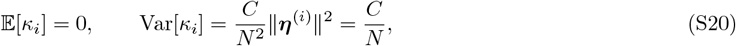

so *κ*_*i*_ ~ *O*(*N*^−1*/*2^) trivially vanishes as *N* → ∞. The self-consistent definition is to replace the 1*/N* normalization with 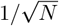:

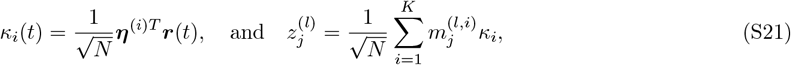

which gives Var[*κ*_*i*_] = *C* = *O*(1) under the GL(*K*) normalization ***η***^(*i*)*T*^ ***η***^(*j*)^ = *Nδ*_*ij*_ that respects 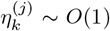 for *k* = 1, …, *N*. Self-consistency of the currents follows immediately:

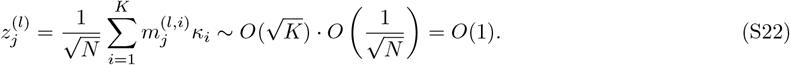

Now, using the formulation of Eq. (S13), consider the latent noise under the 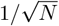 normalization:

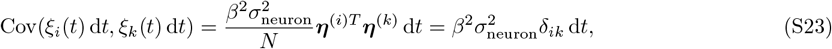

where the diffusion coefficient of *ξ*_*i*_ is *βσ*_neuron_ = *O*(1), independent of *N*. Overall, the *O*(*N*^−1*/*2^) noise suppression of the mean-field limit is entirely absent: in the extensive limit, each latent variable carries *O*(1) noise regardless of population size, and increases in *β* ≫ 1 can no longer be tolerated by increasing *N*. Here, we briefly note that most contemporary works focus primarily on low-rank and *O*(*N*^−1^) scaling of latent variables [59, 60, 61, 62, 63, 94, 95], making it an interesting future direction to study computation in this *O*(*N*^−0.5^) scaling regime [51, 96].

### S2.3 Derivation of the noise covariance matrix

In this appendix, we derive Eq. (6) and thereby prove Proposition 2.

#### S2.3.1. Derivation of Eq. (5)

We apply the integrating factor *e*^*t/τ*^ to Eq. (1) and arrive at:

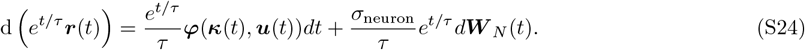

Integrating both sides from 0 to *t* and multiplying through by *e*^−*t/τ*^ yields:

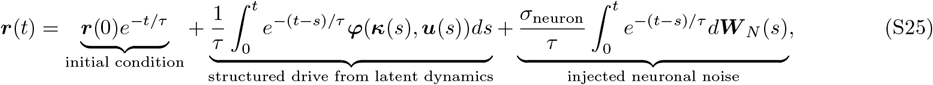

which is Eq. (5) of the main text. Now, the third term is already in its final form, whereas for *t* ≫ *τ*, the initial condition decays to zero and it remains to simplify the structured drive using the separation of neural and latent timescales *β* = *τ*_eff_ */τ* ≫ 1.

#### S2.3.2. Simplification of Eq. (5)

We assume that ***φ*** is bounded and Lipschitz continuous. The latter condition means

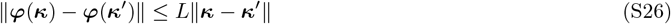

holds for all ***κ, κ***^*′*^ and some *L >* 0. This is satisfied by most standard nonlinearities used in biological neural network models, including tanh, sigmoid, and clipped ReLU. Most importantly, biological constraints require that neural firing rates are bounded and smooth, which justifies both assumptions.

To simplify the structured drive, we add and subtract ***φ***(***κ***(*t*), ***u***(*t*)) inside the integrand:

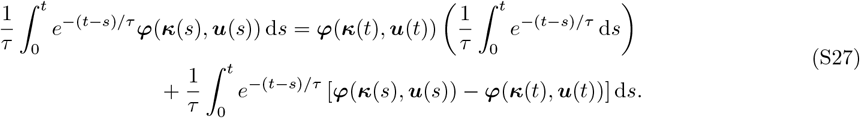

The first term simplifies since

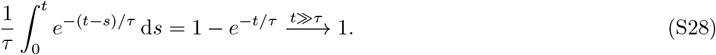

To bound the second term in Eq. (S27), we need to control ∥***κ***(*s*) − ***κ***(*t*)∥. As noted in Proposition 1 and implicit in assumption (iii) of Proposition 2, noise buildup in ***κ***(*t*) within *O*(*τ*) timescales vanishes as *N* → ∞. Since we assume this limit and are interested in scaling with *O*(*β*), we will instead focus on the deterministic part. From Eq. (4b), the deterministic part follows 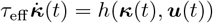, where

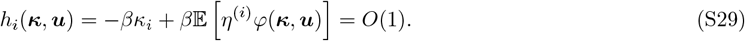

The *O*(1) nature of *h*_*i*_ is by assumption presented in Definition 1, *i.e*., latent variables evolve in timescales *τ*_eff_. Therefore, while the definition of ***h*** contains *β*, by construction the terms are assumed to cancel each other [62]. On the other hand, since *φ* is bounded and *η*^(*i*)^ ~ *O*(1), the second term saturates for large ∥***κ***∥. In this case, the linear term −*βκ*_*i*_ therefore dominates, confining ***κ***(*t*) to a compact set for all *t* ≥ 0. But, this regime also means for large ∥***κ***∥, *h*_*i*_ no longer remains *O*(1), so *κ*(*t*) should be confined in a much smaller ball by assumption.

In short, the notion that ***h*** is *β*-independent requires ***κ*** to remain bounded within a region of validity during normal operation of the network, where *h*_*i*_ remains *O*(1) on observed trajectories. This property in turn means that 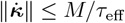 for some *O*(1) constant *M*. For a noiseless network, we therefore have:

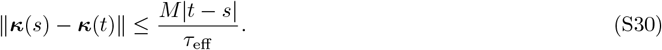

Applying the triangle inequality to move the norm inside the integral, then the Lipschitz condition, and finally substituting the bound on ∥***κ***(*s*) − ***κ***(*t*)∥ gives:

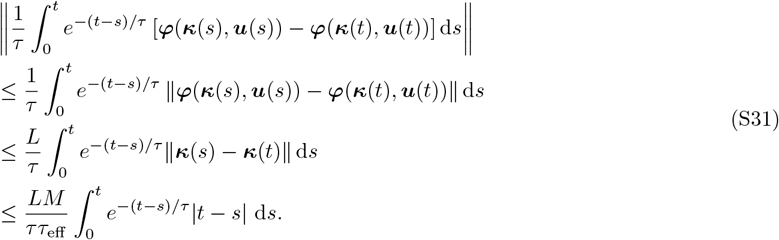

Applying the change of variables *u* = (*t* − *s*)*/τ*, so that |*t* − *s*| = *uτ* and d*s* = *τ* d*u*:

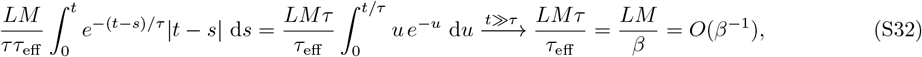

where we used 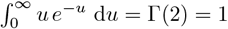. Combining these results gives:

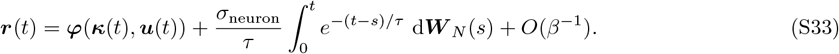

#### S2.3.3. Computing the covariance matrix

From Eq. (S33), the neural activity decomposes as ***r***(*t*) = ***A***(*t*) + ***ε***(*t*) + *O*(*β*^−1^), where we define the structured drive ***A***(*t*) := ***φ***(***κ***(*t*), ***u***(*t*)) and the injected noise 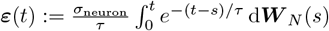. The total population covariance is therefore:

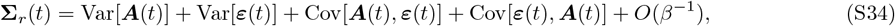

and we compute each term in turn. Since the components *W*_*N,j*_ are independent standard Brownian motions, Cov(*ε*_*j*_ (*t*), *ε*_*k*_ (*t*)) = 0 for *j* ≠ *k*. For *j* = *k*, we apply the Itô isometry, which states that 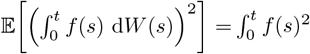 d*s* for any square-integrable deterministic *f* :

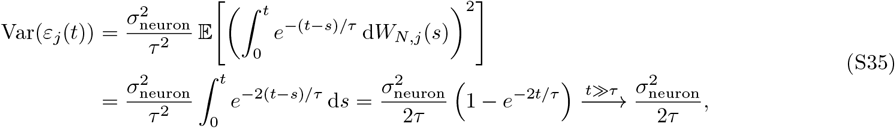

so 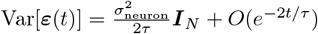.

For the structured drive, following assumption (iii) of Proposition 2, we linearize ***φ***(***κ***(*t*)) around 𝔼[***κ***(*t*)]:

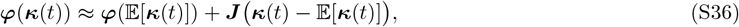

where ***J*** ∈ ℝ^*N ×K*^ has entries *J*_*ji*_ = ∂*φ*_*j*_*/*∂*κ*_*i*_|_𝔼[***κ***(*t*)]_. Since the mean ***φ***(𝔼[***κ***(*t*)]) contributes nothing to the covariance, we obtain Var[***A***(*t*)] = ***J* Σ**_*κ*_(*t*) ***J*** ^*T*^, where **Σ**_*κ*_(*t*) := Var[***κ***(*t*)].

With these, it remains to show that Cov[***A***(*t*), ***ε***(*t*)] = *O*(*β*^−1^). The key observation is that ***ε***(*t*) depends almost entirely on noise increments in the window [*t* − *O*(*τ*), *t*], since the kernel *e*^−(*t*−*s*)*/τ*^ decays on timescale *τ* :

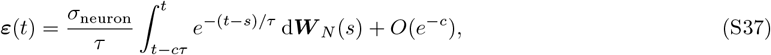

for any *O*(1) constant *c*. The correction is exponential in *c* and does not require any scaling with *β* to vanish (*e.g*., *e*^−5^ *<* 10^−2^), so we do not keep track of it. Since ***κ***(*s*) evolves on the slow timescale *τ*_eff_, the drift bound from the previous section gives ∥***κ***(*t*) − ***κ***(*t* − *cτ*)∥ ≤ *Mcτ/τ*_eff_ = *O*(*β*^−1^), so we decompose:

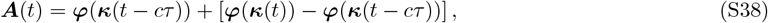

where the second term is *O*(*β*^−1^) by Lipschitz continuity. The first term ***φ***(***κ***(*t* − *cτ*)) depends only on noise increments up to time *t* − *cτ* and is therefore independent of *{*d***W*** _*N*_ (*s*) : *s* ∈ [*t* − *cτ, t*]*}* by the independence of Brownian increments, giving Cov[***φ***(***κ***(*t* − *cτ*)), ***ε***(*t*)] = 0. Combining these results gives Cov[***A***(*t*), ***ε***(*t*)] = *O*(*β*^−1^), and hence:

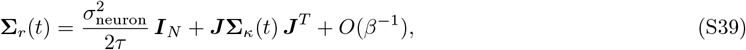

which is Eq. (6) of the main text and proves Proposition 2.

### S2.4 Derivations of the information-limiting noise correlations

In this appendix, we first explicitly identify the differential correlations within our framework. This calculation connects our findings to the mathematics of earlier literature [39, 42] and provides the machinery needed to prove Proposition 3 and Corollary 1.

#### S2.4.1. Normalization of the decoder

Before we start our calculations, we briefly set up the scales for the quantities of interest. Specifically, the linear Fisher information (refer to Eq. (S1)) is independent of ∥***w***∥. Therefore, throughout this section, we choose ***w***^*T*^ 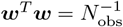 so that individual weights scale as 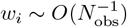, ensuring the decoded signal 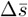 and the elements of the local coding vector 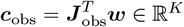 remain both *O*(1) as *N*_obs_ grows. Subsequently, this scaling also ensures that 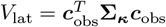 remains *O*(1), *i.e*., independent of *N*_obs_.

#### S2.4.2. Non-zero latent noise in useful decoders

To start with, consider a separation of signal in neural activities **Δ***µ* across two distinct cues. It is trivial to show that for any useful decoder 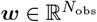, the overlap ***w***^*T*^ **Δ*µ*** ≠0. Otherwise, the Fisher information becomes exactly zero following Eq. (S1). Therefore, any decoder ***w*** that is capable of discriminating the two stimuli cannot be orthogonal to **Δ***µ*.

Now, if two cues (*s*_1*/*2_) at distinct trials were represented closely in the latent subspace, *e.g*., with a small perturbative distance ***κ***_2_ − ***κ***_1_ := *δ****κ***, then we have 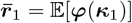 and 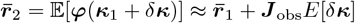]. This means that **Δ*µ*** = *δ****r*** = ***J*** _obs_𝔼[*δ****κ***] ∈ col(***J*** _obs_), where col(***J*** _obs_) refers to the column subspace of 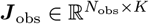. This means that ***w*** has non-zero overlap within col(***J*** _obs_), and therefore 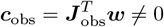 for any useful decoder ***w***, making the contribution of a latent noise term non-zero for such decoders. In other words, this decoder is guaranteed to have a non-zero overlap with col(***J*** _obs_). By Eq. (6), this decoder is also subject to non-vanishing correlated noise. Next, we show that this correlated noise contains differential correlations and therefore limits information.

#### S2.4.3. Differential correlations in the noise covariance matrix

Recall from Eq. (6) that the noise covariance matrix for *N*_obs_ neurons can be written as:

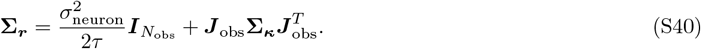

Does this matrix contain contributions of the form **Δ*µ*Δ*µ***^*T*^ with a non-vanishing coefficient? To answer this question for a general matrix ***A***, we perform the following test. Let us write it as:

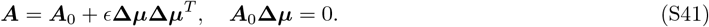

From this, we obtain:

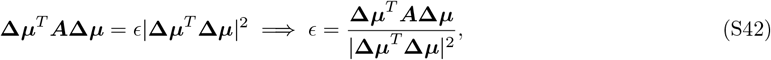

which should remain *ϵ >* 0 as *N*_obs_ → ∞ for differential correlations to exist and limit the Fisher information [39]. Let us consider each term in Eq. (S40) independently.

##### Independent noise injection does not limit information

The first term contains the identity matrix and gives:

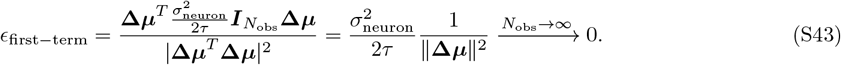

The limit comes from the fact that Δ*µ*_*i*_ ~ *O*(1) for all *i* such that 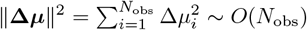. Hence, this term does not have non-vanishing differential correlations and does not limit the information.

##### Accumulated noise in latent variables limits information

Now, consider the second term 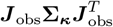, which originates from the noise accumulated in the latent variables. Defining 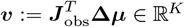 as the projection of the signal direction onto the latent subspace, we obtain:

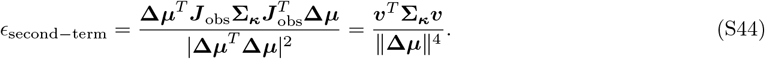

To determine whether this vanishes as *N*_obs_ → ∞, we track the scaling of the numerator and denominator separately. Each entry of 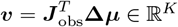 is an inner product of a column of ***J*** _obs_ with **Δ*µ***, which is a sum of *N*_obs_ terms of *O*(1), giving *v*_*i*_ ~ *O*(*N*_obs_). Since the accumulated noise **Σ**_***κ***_ does not depend on the observed number of neurons *N*_obs_, the numerator scales as 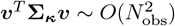. From our calculations above, the denominator similarly scales as 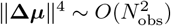. These two scalings cancel exactly, leaving:

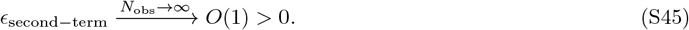

Hence, unlike the first term, *ϵ*_second−term_ remains strictly positive as *N*_obs_ → ∞, confirming that 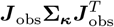 contains genuine differential correlations that limit the extractable Fisher information.

#### S2.4.4. Deriving the saturation curve

Finally, we explicitly show that the accumulated latent noise, which contains the differential correlations, limits the extractable Fisher information. To do so, we first recast Eq. (S1) into a more analytically useful format.

##### The projected discriminability index

To begin with, we claim that Eq. (S1) can be rewritten as:

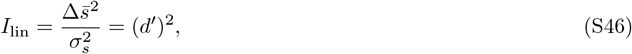

where we defined 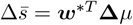 as the projected mean separation with *δ****r*** = **Δ***µ* and 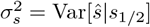 is the noise in the decoder dimension. Its value is given as in Eq. (7) when *s* = *s*_1*/*2_ is fixed, which is equivalent for both classes by assumption. *d*^*′*^ is commonly referred to as the discriminability index [40], which is computed on the projected decoder scores in Eq. (S46).

We can prove this claim by writing the optimal decoder explicitly as 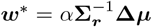 for some scalar *α*, and computing the numerator and denominator of Eq. (S46) separately. For the numerator:

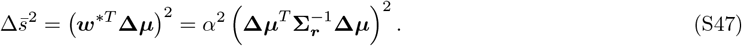

For the denominator, we use 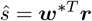 and the definition of **Σ**_***r***_ = Var[***r***|*s*_1*/*2_]:

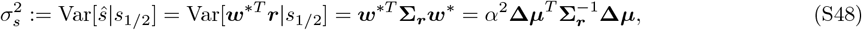

where we used 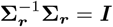 in the last step. Taking the ratio, the *α*^2^ cancels exactly:

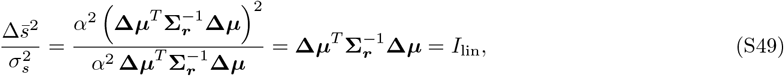

which recovers the original formulation in Eq. (S1). Here, we note that this equation is equivalent to the Fisher information when ***w*** is parallel to the optimal decoder ***w***^∗^. Throughout this work, we refer to the information extractable by a decoder ***w*** to mean this quantity.

##### Information saturation

Recall that, based on our scaling of ***w***, the numerator in Eq. (S46) does not scale with *N*_obs_. Therefore, the scaling of the Fisher information depends on the variance of the linear decoder *ŝ*(*t*) = ***w***^*T*^ ***r***(*t*), which is computed via Eq. (6) as:

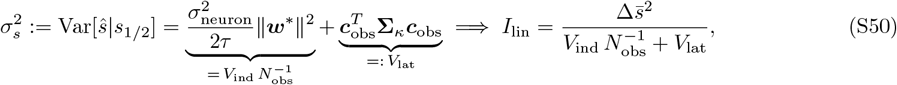

The first term vanishes as *N*_obs_ → ∞, reflecting how pooling over more neurons averages out independent noise. The second term persists regardless of *N*_obs_, since it originates from fluctuations in ***κ*** shared across all neurons through the embedding Jacobian ***J*** _obs_. As we have shown above, this noise term contains differential correlations [39], previously attributed to shared input noise [79]. Here, we have shown that structured synaptic connectivity supporting latent variables produces the same effect by accumulating injected iid noise within *V*_lat_.

#### S2.4.5. Proofs of Proposition 3 and Corollary 1

We now state both proofs explicitly:

**Proof of Proposition 3**

Statement 1 (variance decomposition, Eq. (7)) follows directly from substituting Eq. (6) into the variance of *ŝ*(*t*) = ***w***^*T*^ ***r***(*t*) under the normalization 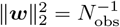, as derived in Eq. (S50). Statement 2 (Fisher information saturation, Eq. (8)) follows from rewriting *I*_lin_ as the discriminability index 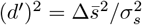 and substituting the decomposition from Statement 1 assuming an optimal decoder 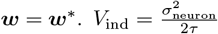 does not depend on *N*_obs_ and the assertion that *V*_lat_ does not scale with *N*_obs_ follows from the normalization established in Appendix S2.4.1. This concludes the proof.

**Proof of Corollary 1**

Any decoder with 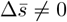 must have ***w***^*T*^ **Δ*µ*** ≠ 0. Since **Δ*µ*** ∈ col(***J*** _obs_) from Appendix S2.4.2, this requires 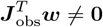 and gives *V*_lat_ *>* 0 as long as **Σ**_*κ*_ is full-rank. Since noise accumulates randomly in latent dimensions via Eq. (4b), this is true by construction. Appendix S2.4.3 then shows that 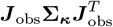 contains genuine differential correlations with a non-vanishing coefficient as *N*_obs_ → ∞, which leads to saturation of information for any decoder ***w*** for which 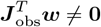. Hence, any useful decoder has a saturating Fisher information, concluding the proof.

### S2.5 Accumulation of latent noise over time

In this section, we prove Proposition 4, which we restate below for completeness:

##### Proposition

(Latent noise accumulation). Consider a latent dynamical system as in Eq. (4b) and assume that ***h*** is smooth, with ***h*** itself and all its derivatives bounded by *O*(1) constants independently of *N* and *β* along any trajectory. Then, for *t* ~ *O*(*τ*_eff_), all eigenvalues of the latent noise covariance scale as:

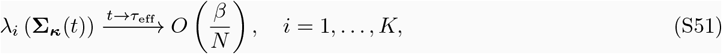

where *λ*_*i*_(**Σ**_***κ***_) denotes the *i*-th eigenvalue of the latent noise covariance matrix **Σ**_*κ*_.

We prove Proposition 4 by linearizing the latent dynamics around the deterministic mean trajectory and bounding the resulting covariance via the Itô isometry. We then verify the bound in three illustrative dynamical regimes (a perfect integrator, a repeller, and an attractive fixed point), where the calculation simplifies and admits exact expressions due to the linearity of latent dynamics [44]. Finally, we combine Propositions 3 and 4 to prove Corollary 2.

#### S2.5.1. Linearization around the deterministic mean trajectory

To start, we rewrite Eq. (4b) as:

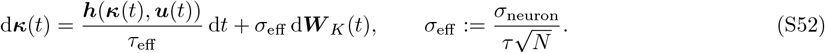

Let 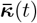 denote the deterministic mean trajectory induced by Eq. (S52) in the noiseless setting:

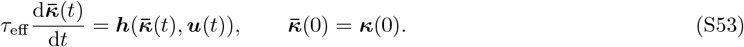

Since ***h*** is bounded by assumption, 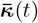 is well-defined for all *t* ≥ 0 and remains in a compact set. Define the fluctuations as 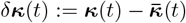. Subtracting the noiseless ODE from Eq. (S52) and Taylor expanding ***h*** around 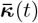 gives:

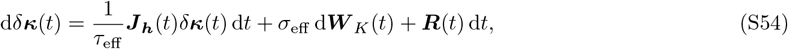

where 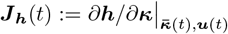 is the Jacobian along the deterministic trajectory and is therefore a fixed function of time, and ***R***(*t*) collects all higher-order Taylor terms. The following lemma justifies dropping ***R***(*t*) in the regime *β* ≪ *N* :

###### Lemma S2

(Self-consistency of the linearization). *Consider the SDE in Eq*. (S54) *under the boundedness assumption on* ***h*** *and its derivatives. At t* ~ *τ*_eff_, *the linearized solution gives* 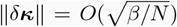, *and the remainder* ***R***(*t*) *is smaller than both the linear drift and the diffusion in the regime β* ≪ *N*.

*Proof*. Dropping ***R***(*t*) entirely yields a linear SDE driven by Brownian noise. The Itô isometry gives 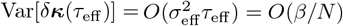, so 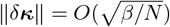. By the boundedness of all derivatives of ***h***, the remainder is bounded by ∥***R***(*t*)∥ = *O*(∥*δ****κ***∥^2^*/τ*_eff_). Comparing the contributions of all three terms in Eq. (S54) to *δ****κ*** over a duration of *τ*_eff_ :

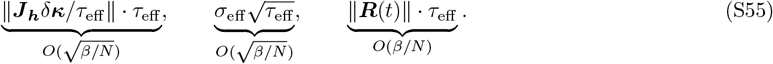

The remainder is smaller than both the linear drift and the diffusion by a factor of 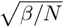, which vanishes in the regime *β* ≪ *N*. Dropping ***R***(*t*) is therefore self-consistent.

While Lemma S2 establishes the trace-norm scaling needed for self-consistency, the eigenvalue-by-eigenvalue scaling claimed in Proposition 4 requires the more detailed analysis below. With ***R***(*t*) controlled, we now solve the linearized SDE around the mean trajectory.

#### S2.5.2. Solving the linearized SDE

Dropping the higher-order term ***R***(*t*), Eq. (S54) reduces to a linear SDE with time-varying drift coefficient 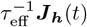. For constant ***J***_***h***_, this would reduce to a multivariate Ornstein-Uhlenbeck process, solvable via eigendecomposition. The time-varying case admits an analogous closed-form solution [45]:

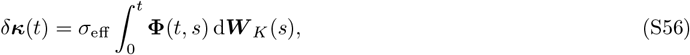

where the state transition matrix **Φ**(*t, s*) ∈ ℝ^*K×K*^ associated with 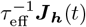 is defined by 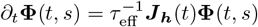 with **Φ**(*s, s*) = ***I***_*K*_. Applying the Itô isometry [45], and using that **Σ**_***κ***_(*t*) = Var[***κ***(*t*)] = Var[*δ****κ***(*t*)] since 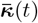 is deterministic:

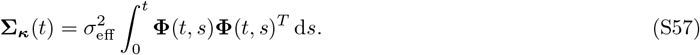

The remaining task is to bound the eigenvalues of this integral.

#### S2.5.3. Bounds on the state transition matrix

By assumption, the largest eigenvalue of 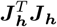 is bounded along any trajectory by some *O*(1) constant *α*^2^, *i.e*., 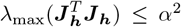. To translate this into a bound on **Φ**(*t, s*), define ***M*** (*t*) := **Φ**(*t, s*)^*T*^ **Φ**(*t, s*), which is symmetric positive semidefinite by construction. Differentiating using the product rule and the defining ODE 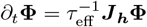:

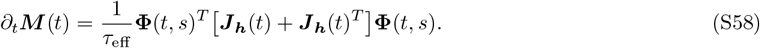

The eigenvalue bound on 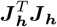 implies 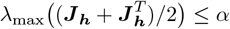, so for any unit vector ***v***:

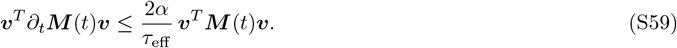

Choosing ***v*** to be the top eigenvector of ***M*** (*t*) at each instant and applying the variational characterization of *λ*_max_ gives the differential inequality *dλ*_max_(***M***)*/dt* ≤ (2*α/τ*_eff_)*λ*_max_(***M***). With initial condition ***M*** (*s*) = ***I***_*K*_, hence *λ*_max_(***M*** (*s*)) = 1, Gronwall’s inequality [45] yields:

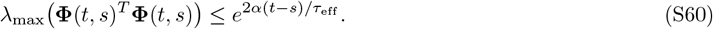

For the lower bound, observe that **Φ**(*t, s*)^−1^ = **Φ**(*s, t*) from the composition property of state transition matrices, and that **Φ**(*s, t*)^*T*^ satisfies an analogous ODE with ***J***_***h***_ replaced by 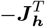. Since the Gronwall argument depends only on 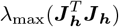, which is invariant under sign flip and transpose, the same upper bound applies:

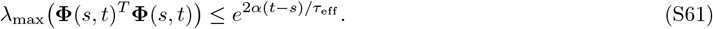

Inverting via the identity *λ*_min_(***A***^*T*^ ***A***) = 1*/λ*_max_(***A***^−1^***A***^−*T*^) for invertible ***A*** yields the lower bound:

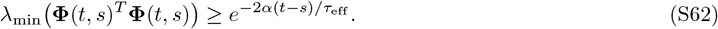

Both bounds hold for all *t* ≥ *s* ≥ 0. For *t, s* ~ *O*(*τ*_eff_) as relevant for our analysis, the exponents are *O*(1) and so are both bounds.

#### S2.5.4. Eigenvalue scaling of the latent noise covariance

We now translate the bounds on **Φ**(*t, s*) into bounds on **Σ**_***κ***_(*t*) via Eq. (S57). The matrices **Φ**^*T*^ **Φ** and **ΦΦ**^*T*^ share the same eigenvalues for any matrix **Φ**, so Eqs. (S60) and (S62) apply directly to **Φ**(*t, s*)**Φ**(*t, s*)^*T*^. For any unit vector ***v***, sandwiching **Σ**_***κ***_(*t*) between the integrated bounds:

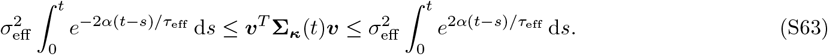

Choosing ***v*** to be each eigenvector of **Σ**_***κ***_(*t*) in turn translates these into bounds on the eigenvalues:

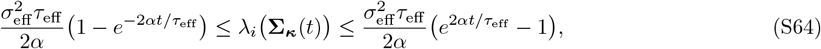

for all *i* = 1, …, *K*. At *t* ~ *τ*_eff_, the bracketed terms are *O*(1), so 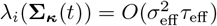. Substituting 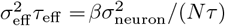:

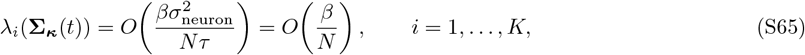

which concludes the proof of Proposition 4.

#### S2.5.5. Three illustrative dynamical regimes

We now verify the general bound in three special cases, where ***J***_***h***_ is constant along the trajectory, and Eq. (S57) admits a closed-form expression as in earlier work [44]. In each case, we recover the *O*(*β/N*) scaling from a direct calculation, providing an independent check of the general result.

##### The case of a perfect integrator

For a perfect integrator, which in the absence of outside input becomes ***h*** = **0**, the SDE in Eq. (S52) reduces to d***κ***(*t*) = *σ*_eff_ d***W*** _*K*_(*t*). This integrates directly to ***κ***(*t*) = ***κ***(0) + *σ*_eff_ ***W*** _*K*_(*t*), and since Var[***W*** _*K*_(*t*)] = *t****I***_*K*_, the latent noise covariance is:

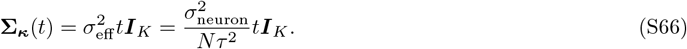

Noise grows linearly in time, unboundedly. At *t* ~ *τ*_eff_ = *βτ*, we obtain 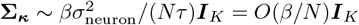.

##### The case of a repeller

For a repeller, the drift term becomes ***h*** ≈ *λ*_+_***κ*** with *λ*_+_ *>* 0, and the SDE in Eq. (S52) becomes d***κ***(*t*) = (*λ*_+_*/τ*_eff_)***κ***(*t*) d*t* + *σ*_eff_ d***W*** _*K*_(*t*). We note that the linear drift ***h*** = *λ*_+_***κ*** is unbounded as ∥***κ***∥ → ∞ and therefore strictly violates the boundedness assumption of Proposition 4. The calculation below should be read as a local linearization valid near the repelling fixed point, with the resulting scaling matching the general bound. Then, applying the integrating factor 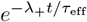, integrating from 0 to *t*, and applying the Itô isometry to the resulting stochastic integral gives:

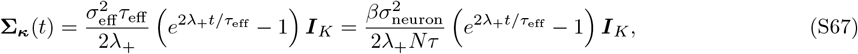

*i.e*., noise grows exponentially in time. At *t* ~ *τ*_eff_, the exponent becomes 2*λ*_+_ = *O*(1), again giving **Σ**_***κ***_ = *O*(*β/N*)***I***_*K*_.

##### The case of an attractive fixed point

For an attractive fixed point, the drift term becomes ***h*** = −*λ*_−_***κ*** with *λ*_−_ *>* 0. Analogous calculations with the integrating factor 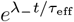 yield:

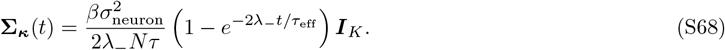

Noise saturates to a stationary value as *t* → ∞, which is expected for an attractive fixed point. At *t* ~ *τ*_eff_, the bracket becomes 1 − *e*^−2*λ*^− = *O*(1), giving **Σ**_***κ***_ = *O*(*β/N*)***I***_*K*_.

In all three regimes, the accumulated latent noise at *t* ~ *τ*_eff_ scales as *O*(*β/N*)***I***_*K*_, independent of the specific form of the drift, in agreement with the general bound of Proposition 4.

#### S2.5.6. Proof of Corollary 2

By Proposition 3, the linear Fisher information saturates at 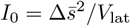, with 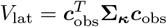 and ∥***c***_obs_∥_2_ = *O*(1) following the normalization in Appendix S2.4.1 and the fact that ***c***_obs_ ∈ ℝ^*K*^ for a fixed *K* ≪ *N*. Substituting the bound from Proposition 4:

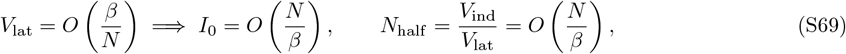

where 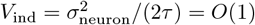 does not scale with *N* or *β*. A non-vanishing *I*_0_ requires *β* ≪ *N*, equivalent to *τ*_eff_ ≪ *Nτ*. The reliable processing window therefore extends to *t* ∈ [0, *O*(*N*)] in units of the neural timescale *τ*, concluding the proof.

## S3 Analyses of Fisher information decay in neocortical recordings

We re-analyze a previously published large-scale calcium imaging dataset [28] in which up to eight neocortical areas were recorded simultaneously in head-fixed mice performing a visual delayed-response task. In each trial, the mouse was shown one of two visual stimuli for 2*s*, followed by a 0.5*s* delay period, after which it reported its choice by licking or withholding a lick. We restrict our analyses to correctly performed trials and to the two binary trial types representing HIT (correct lick) and CR (correct no-lick).

The dataset comprises six animals (L365, L364, L367, L362, L347, L368), each recorded across multiple days totaling to 30 imaging sessions. Neural activity was measured via one-photon calcium imaging and is represented as fluorescence traces sampled at 10*Hz* (Δ*t* = 0.1*s*), yielding 150 time bins per trial spanning −5*s* to +9.9*s* relative to stimulus onset. Neurons were anatomically assigned to one of eight cortical regions: primary visual cortex (V1), lateral visual areas (LV), medial visual areas (MV), posterior parietal cortex (PPC), auditory cortex (A), somatosensory cortex (S), motor cortex (M), and retrosplenial cortex (RSC). An additional “all-areas” condition (ALL) pools every recorded neuron across regions. For any given (area, animal) pair, we require at least 50 simultaneously recorded neurons with no missing time points. Pairs that do not meet this threshold are excluded from all analyses.

### S3.1 Decoding analysis and estimating the Fisher information

#### PLS-LDA pipeline

We follow the decoding practices of the prior work [28]. At each time bin *t*, neural activity is summarized by the matrix 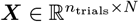, where *N* is the number of recorded neurons. To decode the binary trial identity from ***X***, we use a two-stage dimensionality-reduction and classification pipeline combining Partial Least Squares (PLS) regression and Linear Discriminant Analysis (LDA).

For a given number of PLS components *d*_PLS_ ∈ *{*1, …, 15*}*, each decoding experiment proceeds as follows. Trials are first split into a training set (two-thirds) and a held-out test set (one-third), stratified by trial type. The training set is further divided into two equal halves. PLS regression with *d*_PLS_ components is fit on the first half of the training set with trial identity as the response variable. The resulting model is used to project the second half of the training set into the *d*_PLS_-dimensional PLS subspace, and (with analytical shrinkage of the within-class covariance [97]) is then fit on these projected activations.

The LDA weight vector 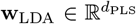 is projected back to the full neural space via **w** = **P**_rot_**w**_LDA_, where 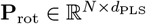 is the matrix of PLS *x*-rotation vectors. The combined decoder **w** is then evaluated on the held-out test set. Decoding performance is quantified by the squared discriminability index *d*^*′*2^, defined as

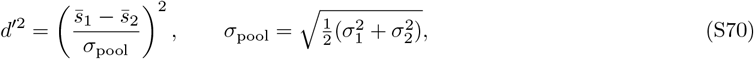

where 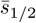 and *σ*_1*/*2_ are the mean and standard deviation of the projected test-set scores *ŝ* = **w**^*T*^ **r***/*∥**w**∥_2_ in each trial class. Under the Gaussian noise assumption, *d*^*′*2^ is the linear Fisher information *I*_lin_ in natural units [40]. Given the near optimality of linear decoders on this dataset [62], we use it as our empirical proxy for *I*_lin_ throughout. This procedure is repeated for 100 independent random train/test splits, and the resulting *d*^*′*2^ values are averaged across splits.

#### Decoding accuracy as a function of PLS dimensionality (Fig. S1)

To assess how many latent dimensions are needed to capture the task-relevant signal, we compute *d*^*′*2^ as a function of *d*_PLS_ for each time bin and then average over bins within three task epochs: the stimulus window (0-2*s*), the delay window (2-2.5*s*), and the response window (2.5-5*s*). For each (area, animal) pair with at least 50 neurons, the cross-split-averaged *d*^*′*2^ is computed separately for each *d*_PLS_ ∈ *{*1, …, 15*}*. Figure S1 shows the cross-animal mean *±* SEM of *d*^*′*2^ as a function of *d*_PLS_ for each cortical area and each task epoch. The curves plateau rapidly and we use *d*_PLS_ = 15 for our later analyses.

#### Temporal decay of Fisher information (Figs. 2 and S2)

We next examine how *d*^*′*2^ evolves over the full trial time course. For each (area, animal) pair, the 100-split average of *d*^*′*2^ is computed at every 100 ms time bin. Figure 2 shows, for each area, the cross-animal mean *d*^*′*2^ together with a *±*1 SD band across animals.

To quantify the decay timescale, we fit a single exponential *d*^*′*2^(*t*) ∝ exp(−*t/τ*_eff_) to each trace in the post-trial window from *t* = 5*s* to *t* = 10*s* (gray shaded region in Fig. 2), which falls after the response period and is presumably uncontaminated by task-evoked transients. The fit is performed by ordinary least-squares linear regression of log *d*^*′*2^(*t*) on *t*; the decay constant is 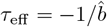, where 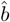 is the estimated slope. For Fig. 2, the fit is applied to the cross-animal mean trace of each area. For Fig. S2, the same fit is applied to each individual animal’s trace within each area separately, and the panel annotation reports the mean *±* SEM of *τ*_eff_ across animals.

### S3.2 Control analyses with single-neuron autocorrelation timescales

A potential confound for the slow post-trial decay of *d*^*′*2^ is that individual calcium transients or slowly varying neurons could artificially inflate the apparent population timescale. We address this concern in two steps.

#### Estimating single-neuron autocorrelation timescales (Fig. S3)

For each neuron, we compute the autocorrelation function (ACF) at lags *k* = 0, 1, …, 49 (i.e., 0 to 4.9*s*). Specifically, for lag *k*, we concatenate the activity from all trials over the reference window [0*s*, 5*s*) and the corresponding shifted window [*k*Δ*t*, 5*s* + *k*Δ*t*), and compute the Pearson correlation coefficient between these two sample vectors.

We estimate each neuron’s autocorrelation timescale using two complementary measures. *(i)* Exponential-fit timescale *τ*_fit_: a weighted log-linear regression of log ACF(*k*) on *k* is performed over lags *k* = 0, …, 10 (0 to 1 s), with weights equal to the ACF value at each lag (which down-weights lags where the ACF is already near zero). The estimated slope 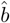 gives 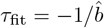 in units of time bins. *(ii)* Half-crossing timescale *τ*_1*/*2_: the first lag at which the ACF drops below 0.5 is linearly interpolated to give a half-life *t*_1*/*2_, from which *τ*_1*/*2_ = *t*_1*/*2_*/* ln 2.

Figure S3**A** shows up to 100 randomly selected ACF traces per cortical area. Figure S3**B** shows the cumulative fraction of neurons whose timescale is at most *t*, computed per animal and averaged across animals as mean *±* SEM. An animal is included in this summary for a given area only if it contributes at least 50 neurons with a finite timescale estimate in that area.

#### Decoding after excluding slowly-autocorrelated neurons (Fig. S4)

To verify that the slow population-level decay of *d*^*′*2^ is not inherited from a small subset of cells with long calcium transients, we repeat the full PLS-LDA decoding analysis after removing neurons whose single-cell autocorrelation timescale exceeds a threshold *τ*_max_. Specifically, a cell is retained only if both *τ*_fit_ and *τ*_1*/*2_ are finite and do not exceed *τ*_max_. We sweep *τ*_max_ ∈ *{*0.5, 0.7, 1.0*}* s, corresponding to caps of 5, 7, and 10 time bins, respectively. For each value of *τ*_max_, the decoding pipeline is run with *d*_PLS_ = 15 PLS components and 100 random splits, and the same post-trial exponential fit (5-10*s*) is applied to the resulting mean *d*^*′*2^ traces. Figure S4 shows that the fitted decay constants *τ*_eff_ remain mostly in the range [2, 3]*s* across all three thresholds and are comparable to those obtained without any cell filtering (Fig.2).

## S4 Testing the latent dimensionality

### S4.1 A covariance-matching test of latent dimensionality

In this section, we explain the process of least-squares covariance matching, which we use to estimate the dimensionality of the latent subspace via Eq. (6):

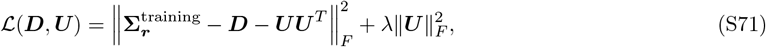

where 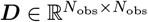 is a non-negative diagonal matrix absorbing the independent noise floor 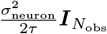 in Eq. (6), 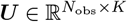 encodes the rank-*K* correlated component ***J* Σ**_***κ***_***J*** ^*T*^, and *λ* ≥ 0 is an *ℓ*_2_ regularization parameter selected by cross-validation. The symmetric factorization ***UU*** ^*T*^ enforces positive semi-definiteness of the low-rank term, consistent with its interpretation as a covariance matrix. The regularization term 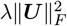 penalizes large loadings in ***U***, preventing the low-rank component from absorbing variance that properly belongs to the independent noise floor ***D***. The optimal *λ* is selected by cross-validation over a logarithmically spaced grid.

#### S4.1.1. Minimization via alternating updates

We minimize by alternating between two closed-form updates. The update for the ***D*** is trivial, which we show first.

For a *k*th step estimate ***U*** ^(*k*)^, start by defining the residual 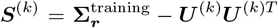. With ***U*** ^(*k*)^ fixed, 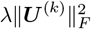 is constant and the objective reduces to minimizing 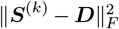 over non-negative diagonal ***D***. Since ***D*** is diagonal, it only enters through the diagonal entries of the residual, so the Frobenius norm decomposes as

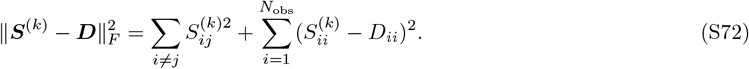

The off-diagonal terms are independent of ***D***, so the optimal update is a pointwise projection onto the non-negative real values:

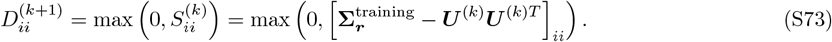

#### S4.1.2. Solving the second subproblem analytically

To solve the second subproblem, with ***D***^(*k*+1)^ fixed, define 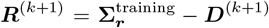, which is symmetric by definition. The subproblem to solve for ***U*** ^(*k*+1)^ is

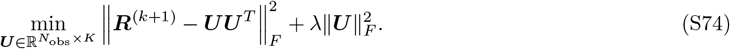

Since ***UU*** ^*T*^ is symmetric positive semi-definite with rank at most *K*, we reparametrize as ***U*** = ***P* Γ**^1*/*2^, where 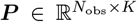 has orthonormal columns (***P*** ^*T*^ ***P*** = ***I***_*K*_) and **Γ** = diag(*γ*_1_, …, *γ*_*K*_) with *γ*_*k*_ ≥ 0. Under this parametrization, ***UU*** ^*T*^ = ***P* Γ*P*** ^*T*^ and 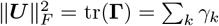, so Eq. (S74) becomes

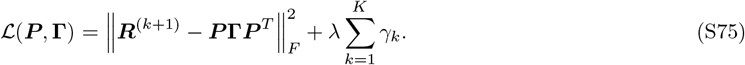

To solve this, we first let ***R***^(*k*+1)^ = ***V MV*** ^*T*^ be the eigendecomposition of ***R***^(*k*+1)^, with eigenvalues 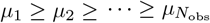 and eigenvectors ***V***. This follows from the observation that ***R***^(*k*+1)^ is a symmetric matrix. Expanding the Frobenius norm via the cyclic property of the trace, we obtain:

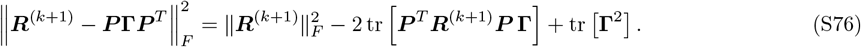

Here, we used tr[(***P* Γ*P*** ^*T*^)^2^]= tr[**Γ**^2^***P*** ^*T*^ ***P]***= tr[**Γ**^2^]. For fixed **Γ**, minimizing over ***P*** requires maximizing 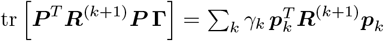. This is achieved when the columns of ***P*** are the top-*K* eigenvectors of ***R***^(*k*+1)^:

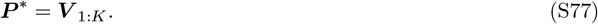

Now, with ***P*** = ***V*** _1:*K*_, the eigendecomposition of ***R***^(*k*+1)^ gives ***P*** ^*T*^ ***R***^(*k*+1)^***P*** = diag(*µ*_1_, …, *µ*_*K*_), so the Frobenius norm decomposes across eigenspaces as

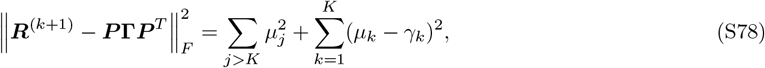

and the full objective reduces to *K* independent scalar problems:

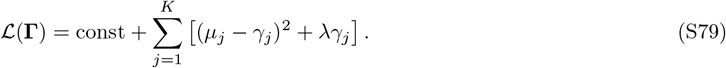

Differentiating with respect to *γ*_*j*_ and applying the non-negativity constraint *γ*_*j*_ ≥ 0, which ensures ***UU*** ^*T*^ = ***P* Γ*P*** ^*T*^ remains positive semi-definite, gives:

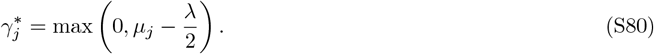

Combining both steps, the optimal low-rank term for the (*k* + 1)th iterate is

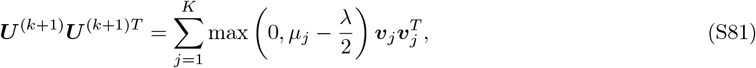

where *µ*_*j*_ and ***v***_*j*_ are the eigenvalues and eigenvectors of 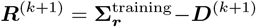. This is a low-rank reconstruction of ***R***^(*k*+1)^, but any dimension whose eigenvalue falls below *λ/*2 is eliminated entirely.

The two alternating updates are iterated to convergence, yielding the final estimates 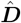 and ***Û***. We have written a custom script to implement this process, which will be publicly available with our codebase upon publication.

#### S4.1.3. Initialization and stopping criteria

The algorithm is initialized by computing the top-*K* eigenvectors of 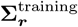, soft-thresholding the corresponding eigenvalues by *λ/*2 to obtain ***U*** ^(0)^, and setting ***D***^(0)^ to the non-negative diagonal residual 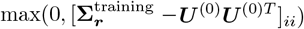. The alternating updates are then run for a maximum of 500 iterations. Convergence is declared when the relative decrease in training loss falls below a tolerance *ϵ* = 10^−6^, *i.e*.,

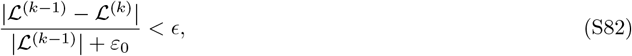

where *ε*_0_ = 10^−12^ prevents division by zero, and at least one full iteration is always performed.

### S4.2 Tests with task-trained artificial neural networks

#### S4.2.1. Training details

##### Flip-flop task

The *L*-bit flip-flop task requires a network to independently maintain the most recent *±*1 input for each of *L* channels. At each timestep, the input ***u***(*t*) ∈ ℝ^*L*^ for each channel is either 0 (no event) or *±*1 (a flip event drawn uniformly from *{*−1, +1*}*). Flip candidates are drawn independently at each timestep with probability 0.2, and any candidate occurring within four timesteps of a previously accepted flip is suppressed. The target output for each channel is the value of the most recent flip event on that channel, *i.e*., each channel independently maintains a persistent binary memory. Training sequences have length *T* = 100 and are done in batches of *B* = 100.

##### Vanilla RNNs

The first class of networks consists of vanilla RNNs whose hidden state evolves as

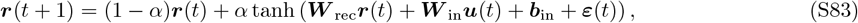

with leak rate *α* = 0.5, a full *N × N* recurrent weight matrix ***W*** _rec_, an input projection ***W*** _in_ ∈ ℝ^*N ×L*^ with bias ***b***_in_ ∈ ℝ^*N*^, and a linear readout ***ô***(*t*) = ***W*** _out_***r***(*t*) + ***b***_out_ with ***W*** _out_ ∈ ℝ^*L×N*^ and bias ***b***_out_ ∈ ℝ^*L*^. Here ***ε***(*t*) ~ 𝒩 (0, *σ*^2^***I***_*N*_) is iid Gaussian noise injected additively into the pre-activation argument of the nonlinearity at each discrete timestep, where *σ* is the noise level as in Fig. S5.

##### LSTMs

The second class consists of standard LSTMs whose gate updates and state dynamics are given by

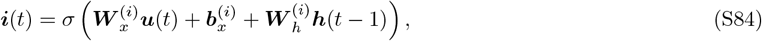

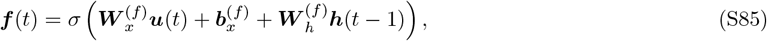

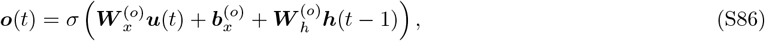

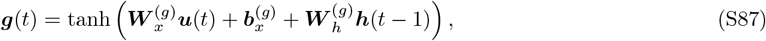

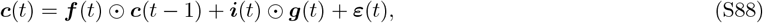

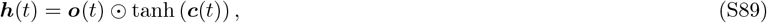

where *σ*(·) denotes the sigmoid function, ⊙ denotes elementwise multiplication, with the input-to-gate matrices 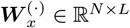, biases 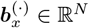, and the hidden-to-gate matrices 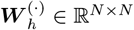. The noise ***ε***(*t*) ~ 𝒩 (0, *σ*^2^***I***_*N*_) is injected additively into the cell state ***c***(*t*) after the standard gating update. The readout is ***ô***(*t*) = ***W*** _out_***h***(*t*) + ***b***_out_ with ***W*** _out_ ∈ ℝ^*L×N*^ and bias ***b***_out_ ∈ ℝ^*L*^. Both architectures use hidden dimension *N* = 100.

##### Training

All weight matrices were initialized with Xavier uniform initialization. Networks were trained without injected noise (*σ* = 0). Networks were optimized with Adam (learning rate 10^−3^) for 2000 epochs using mean-squared error loss, with gradient clipping at a global norm of 1.0. We trained 20 independent random seeds per *L* and architecture, yielding 200 trained networks in total.

#### S4.2.2. Estimating the noise covariance matrix

##### Extracting the noise fluctuations

To estimate the noise fluctuations, we loaded each trained network, fixed its weights, and recorded hidden-state activity along long flip-flop sequences of length *T* = 10,000 with a reduced flip probability of *p*_flip_ = 0.02 (before gap enforcement), injecting iid Gaussian noise at each timestep with standard deviation *σ* ∈ *{*10^−3^, 10^−2^, 10^−1^*}* as described above. We discarded the first 100 timesteps to eliminate transient initialization effects. To isolate noise fluctuations from the signal, we subtracted per-state conditional means from the hidden-state trajectories. The latent state was encoded as a scalar label 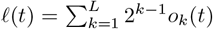, where *o*_*k*_(*t*) ∈ *{*−1, +1*}* is the *k*-th channel output at time *t*. For each unique value of *ℓ*, we subtracted the mean hidden state across all timesteps sharing that label, yielding mean-centered residuals *δ****r***(*t*) that reflect noise fluctuations around the signal.

##### Fitting the noise covariance matrix

We estimated the empirical noise covariance from a contiguous 80*/*20 train/test split of the mean-centered time series, with a random circular shift applied before splitting to decorrelate the boundary. We computed the held-in **Σ**_train_ and held-out **Σ**_test_ covariance matrices. We then fit a diagonal-plus-rank-*K* model 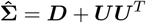 to **Σ**_train_ by alternating minimization as described in Appendix S4.1, sweeping over *K* ∈ *{*1, …, 10*}* and regularization strengths *λ* on a logarithmic grid of ten values spanning [10^−6^, 10^0^]. For each (*K, λ*) pair, we evaluated generalization by computing the Pearson correlation between the off-diagonal entries of the estimated 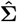 and **Σ**_test_. The best *λ* was selected by maximizing this test correlation. We repeated this procedure across *n*_exp_ = 10 independent long sequences per network and averaged results across sequences and the 20 trained seeds. Results are presented in Fig. S5.

### S4.3 Applications to neocortical recordings

#### S4.3.1. Covariance matching

We applied the covariance-matching framework to the same large-scale calcium imaging dataset described in Appendix S3, consisting of six animals (L347, L362, L364, L365, L367, L368) recorded across multiple days. For each animal and each cortical area, we required at least 301 simultaneously recorded neurons and area-animal pairs not meeting this threshold were excluded. As a result, the motor cortex did not meet this threshold for any animal and was excluded from this analysis. Hence, neural activity was analyzed across seven cortical regions: V1, LV, MV, PPC, A, S, RSC as described in the caption of Fig. 2. Plus, as before, we also used all neurons pooled across areas (ALL). This analysis was done on all trials, correctly or incorrectly performed.

##### Mean centering

Neural activity was mean-centered separately for each recording day before pooling across days. Within each day, we subtracted the per-trial-type conditional mean from each neuron’s activity and for each time *t* into the trial. We pooled data from all four trial types (HIT, CR, and the two incorrect trial types) to estimate the trial-to-trial variability in neural activities. These were then concatenated across all days, yielding a single array of shape (*n*_trials_, *T, N*) per animal and area.

##### Time windows

We analyzed ten non-overlapping time windows of five bins each (0.5 s), spanning bins 50 to 100 of the trial (corresponding to 0 s to 5 s relative to stimulus onset, covering the stimulus, delay, and response periods). Within each window, trials and time bins were flattened along the time axis before computing covariance matrices.

##### Covariance estimation and factorization

For each time window, we performed *n*_exp_ = 20 independent random 80*/*20 train/test splits at the trial level. For each split, we computed the held-in covariance matrix **Σ**_train_ and held-out covariance matrix **Σ**_test_. We then fit a diagonal-plus-rank-*K* model 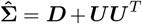 to **Σ**_train_ by alternating minimization as described in Appendix S4.1, sweeping over *K* ∈ *{*1, 2, 3, 4, 5, 6, 7, 8, 9, 10, 15, 20, 30, 50, 100*}* and regularization strengths *λ* on a logarithmic grid of ten values spanning [10^−2^, 10^2^]. For each (*K, λ*) pair, generalization was evaluated by the Pearson correlation between the off-diagonal entries of 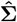 and **Σ**_test_. The best *λ* was selected independently for each fitting instance by maximizing this test correlation.

##### Dimensionality estimate

To extract a scalar dimensionality estimate from the test correlation curve as a function of *K*, we fit a saturating exponential *A*(1 − *e*^−*K/τ*^) + *c* to each curve after replacing the values after the peak with the peak value to handle non-monotone curves arising from over-regularization at large *K*. The dimensionality estimate is defined as *d*_CM_ = 3*τ*, which corresponds to the rank at which the saturating exponential has reached 95% of its asymptotic value. For a given animal and brain region, the estimated test correlations were averaged across the 20 splits and are presented in Figs. 3 and S7.

#### S4.3.2. Participation ratio analysis

##### Participation ratio

Proposition 2 predicts that the noise covariance matrix takes a diagonal-plus-low-rank form. An important consequence is that if the diagonal independent-noise term dominates, the full covariance matrix will appear high-dimensional by any linear measure. Classical dimensionality measures applied to the full covariance matrix therefore serve as a baseline that is expected to yield high dimensionality, irrespective of the true rank of the correlated component. To verify this prediction and introduce an interpretable baseline for the covariance-matching approach, we computed the participation ratio (PR) of both the full and off-diagonal noise covariance matrices. The PR is defined as

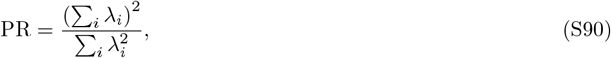

where *λ*_*i*_ are the eigenvalues of the covariance matrix, and provides a continuous estimate of the effective number of dimensions that carry variance.

##### Full and low-rank PR calculations

We computed the PR under two conditions. In the first, we applied the PR to the full sample covariance matrix **Σ**_***r***_, including its diagonal. In the second, we set the diagonal entries of **Σ**_***r***_ to zero before computing the eigendecomposition, isolating the shared variance structure and suppressing the contribution of independent neuronal noise. We then used the positive eigenvalues to compute the participation ratio. The mean-centering, day-concatenation, time-window, and neuron-inclusion procedures were identical to those described in the covariance-matching analysis above. The PR was computed separately for each time window and each animal, and results are presented in Figs. 3 and S6.

#### S4.3.3. Factor analysis

##### Fitting of the factor analysis

As a second baseline for dimensionality estimation, we applied standard factor analysis (FA) implemented in scikit-learn [98]. FA assumes a generative model in which observed activity is driven by *K* latent Gaussian factors plus independent neuron-specific noise, and unlike the covariance-matching approach it requires an explicit Gaussian distributional assumption on the structured noise component. The data preparation, trial-type selection, neuron-inclusion threshold, and time-window definitions were identical to those described in the covariance-matching analysis above. For each time window and each of *n*_exp_ = 20 independent random 80*/*20 train/test splits at the trial level, we fit FA models with *K* ∈ *{*1, 2, 3, 4, 5, 6, 7, 8, 9, 10, 12, 15, 18, 20, 25, 30, 40, 50, 75, 100*}* latent factors to the held-in data and evaluated generalization by computing the per-sample log-likelihood on the held-out data. The median held-out log-likelihood across test samples was used as the generalization metric, to remain robust to outlier time points.

##### Dimensionality estimation

To extract a scalar dimensionality estimate, the log-likelihood curves as a function of *K* were normalized within each fitting instance by subtracting the minimum and dividing by the absolute maximum, placing all curves on a common [0, 1] scale. We then fit a saturating exponential *A*(1 − *e*^−*K/τ*^) + *c* to each normalized curve using the same plateau-filling and fitting procedure described for the covariance-matching analysis. The dimensionality estimate is defined as *d*_FA_ = 3*τ*, consistent with the 95% saturation criterion used throughout. Results were averaged across the 20 splits and are reported in Figs 3 and S8.

## S5 Behavioral training and delay-generalization analysis

Behavioral experiments were performed in male C57BL/6J mice, which began training at 11-12 weeks of age. Mice were water restricted and received 1ml water per day, with no additional free access to water outside the training schedule. Training was conducted during the animals’ active circadian phase, and each mouse was trained at a fixed time of day across sessions.

### Task details

Mice were trained on a visual two-alternative forced choice (2AFC) delayed discrimination task (Fig. S9). In each trial, mice were presented with a visual sample cue (blue or yellow), followed by a delay period and a response epoch during which lick ports became available. Cue–response contingencies were fixed throughout training: the blue cue instructed a left lick, whereas the yellow cue instructed a right lick. A correct response was defined as licking to the cue-associated port during the response window.

### Progressive delay experiment

For the progressive delay-learning analysis shown in Figs. 4 and S9, 9 male C57BL/6J mice were trained in two stages. In Stage 1, mice were trained at 0 s delay. Once performance exceeded 70% correct in a session, mice advanced to Stage 2, in which the delay duration was increased in 0.25s increments. Mice remained at a given delay until the performance criterion was reached, after which the next delay level was introduced. This procedure continued until animals achieved stable performance at a 3s delay. Behavioral sessions typically consisted of approximately 150 trials. For sessions used to assess whether animals reached criterion, trials showing clear disengagement were excluded before recalculating session accuracy. Specifically, if performance at the beginning of a session was markedly lower than the overall session average, the first 20 trials were removed. In addition, once a sustained sequence of no-lick trials appeared near the end of a session, the subsequent trials were excluded. After this trimming procedure, the sessions still contained at least 100 trials, which were used to determine whether the 70% performance criterion had been reached.

For each mouse and each delay level, the trials-to-criterion value was defined as the cumulative number of trials required from the first session at that delay until the first session in which performance exceeded 70% correct, provided that the session contained at least 100 completed trials. If a mouse did not reach criterion at a given delay in a given session, training at that delay continued across subsequent sessions until criterion was reached. Thus, trials-to-criterion values were accumulated across sessions and were obtained for every mouse at every delay level included in the analysis.

To quantify learning burden across delay regimes, delay levels were grouped into three bins: short (0–0.75 s), target (1–1.25 s), and long (1.5–3 s). The intermediate bin was chosen to bracket the theory-motivated transition near ~1s while respecting the discrete 0.25s training increments used in the task. For each mouse, trials-to-criterion values were first averaged within each bin, and these mouse-level averages were then compared across bins. The primary statistical analysis used two-sided paired t-tests. To assess robustness of the critical delay-bin comparisons, two-sided Wilcoxon signed-rank tests were additionally performed for the comparisons of the target interval against the short- and long-delay bins.

### Generalization to longer delays

This was assessed by splitting mice from the progressive delay experiment into two groups, results shown in Fig. S11. In the constant-delay 6s generalization test, 5 male C57BL/6J mice that had been trained to stable performance at 3s delay were tested in a session consisting of a block of 120 trials at 3s delay, immediately followed by 60 trials at 6s delay. Performance was quantified as percent correct in each block, and two-sided paired t-tests were used to compare the 3s and 6s conditions at the mouse level. In the random-delay 6 s generalization test, 4 male C57BL/6J mice were tested in a single session of similar length, in which 6s delay trials comprised 20% of trials and were randomly interleaved with standard 3s delay trials. Performance was quantified separately for 3s and 6s trials within the same session, and two-sided paired t-tests were used to compare percent correct between the two delay conditions at the mouse level.

## S6 Supplementary Figures

**Figure S1.**
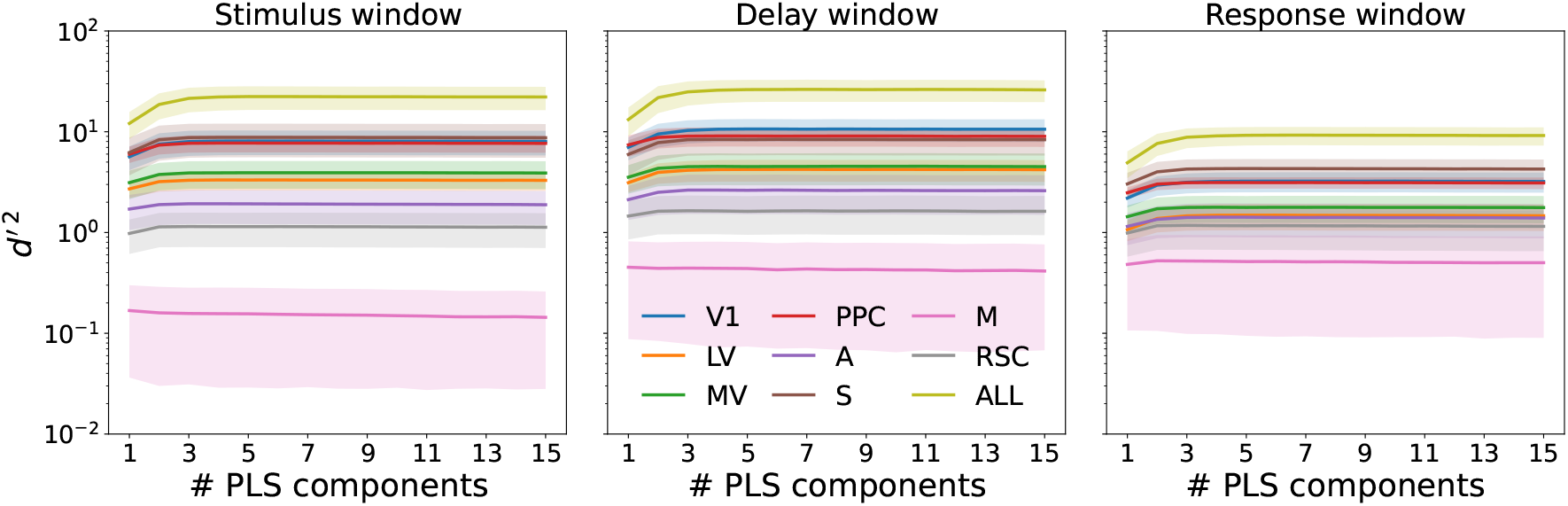
Decoding accuracy as a function of PLS dimensionality across cortical areas and task epochs. Fisher information (*d*^*′* 2^) of correct trial identity (HIT vs CR) decoded from population activity, plotted against the number of PLS components retained prior to LDA. Decoders were trained and tested on held-out trials within each animal (*n* = 6) for each time point (100ms bins). Decoder *d*^*′* 2^ values were averaged across 100 random splits and across timepoints within the stimulus window (left), delay window (middle), and response window (right). Curves show the cross-animal mean for each cortical area: primary visual cortex (V1), lateral visual areas (LV), medial visual areas (MV), posterior parietal cortex (PPC), auditory cortex (A), somatosensory cortex (S), motor cortex (M), retrosplenial cortex (RSC), and all areas pooled (ALL). Analysis was restricted to area-animal combinations with at least 50 simultaneously recorded neurons. Shaded bands denote SEM across available number of animals.

**Figure S2.**
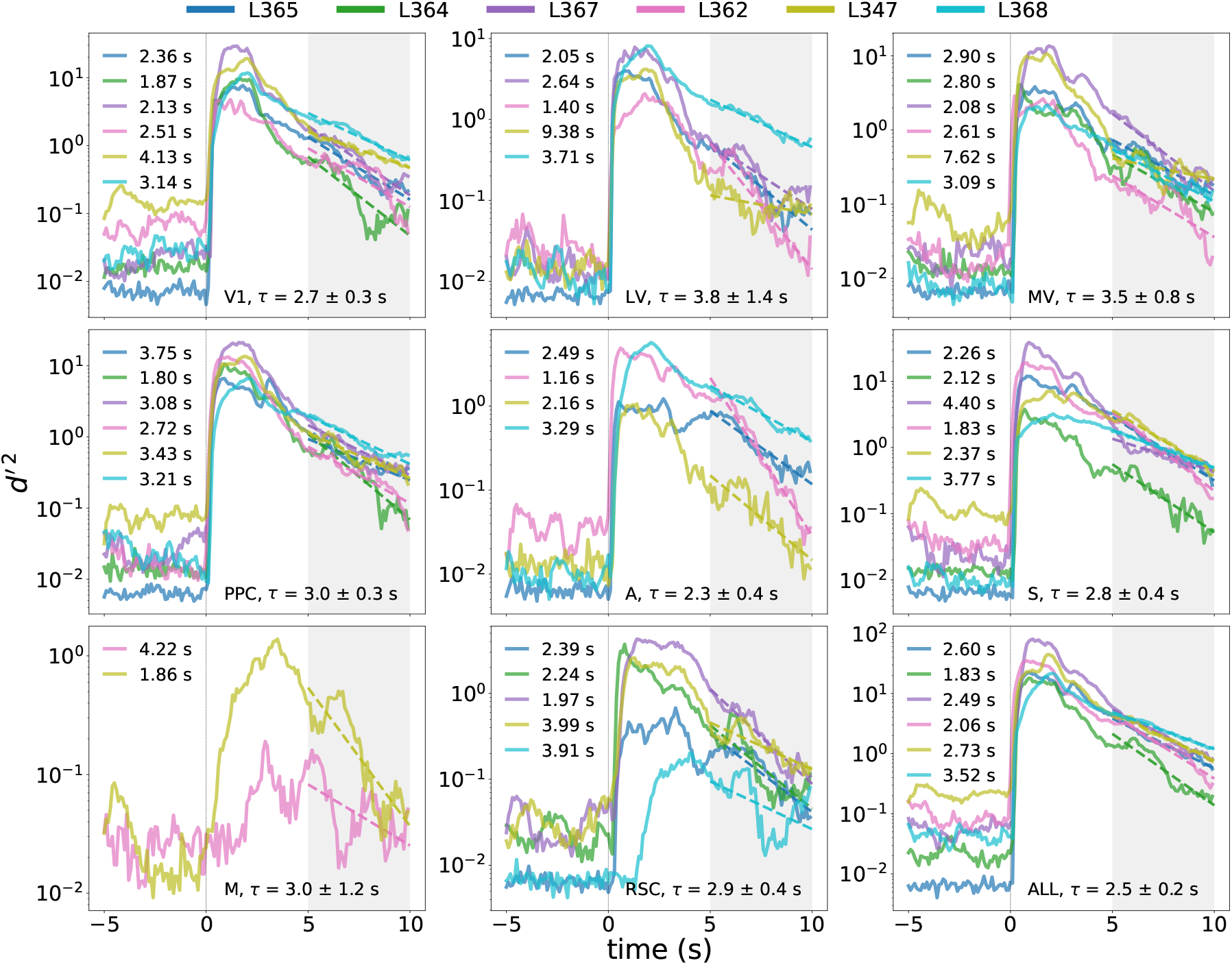
Decay of decoding accuracy after the trial conclusion. Time course of the Fisher information (*d*^*′* 2^), decoded from population activity using PLS regression with 15 components followed by LDA, and averaged across 100 random train/test splits per animal. Each panel shows one cortical area. Time *t* = 0 s marks stimulus onset (vertical line); the shaded gray region (*t* = 5 s to *t* = 10 s) indicates the after-trial time window over which the exponential decay was fit. Solid traces show individual animals (L365, L364, L367, L362, L347, L368; see [28]), restricted to area-animal combinations with at least 50 simultaneously recorded neurons. Dashed lines show per-animal exponential fits of the form *d*^*′* 2^(*t*) ∝ exp(−*t/τ*), obtained by a linear regression on log *d*^*′* 2^. Per-animal time constants *τ* are listed in each panel, whereas the area-level summary (bottom of each panel) reports the across-animal mean *±* sem of *τ*.

**Figure S3.**
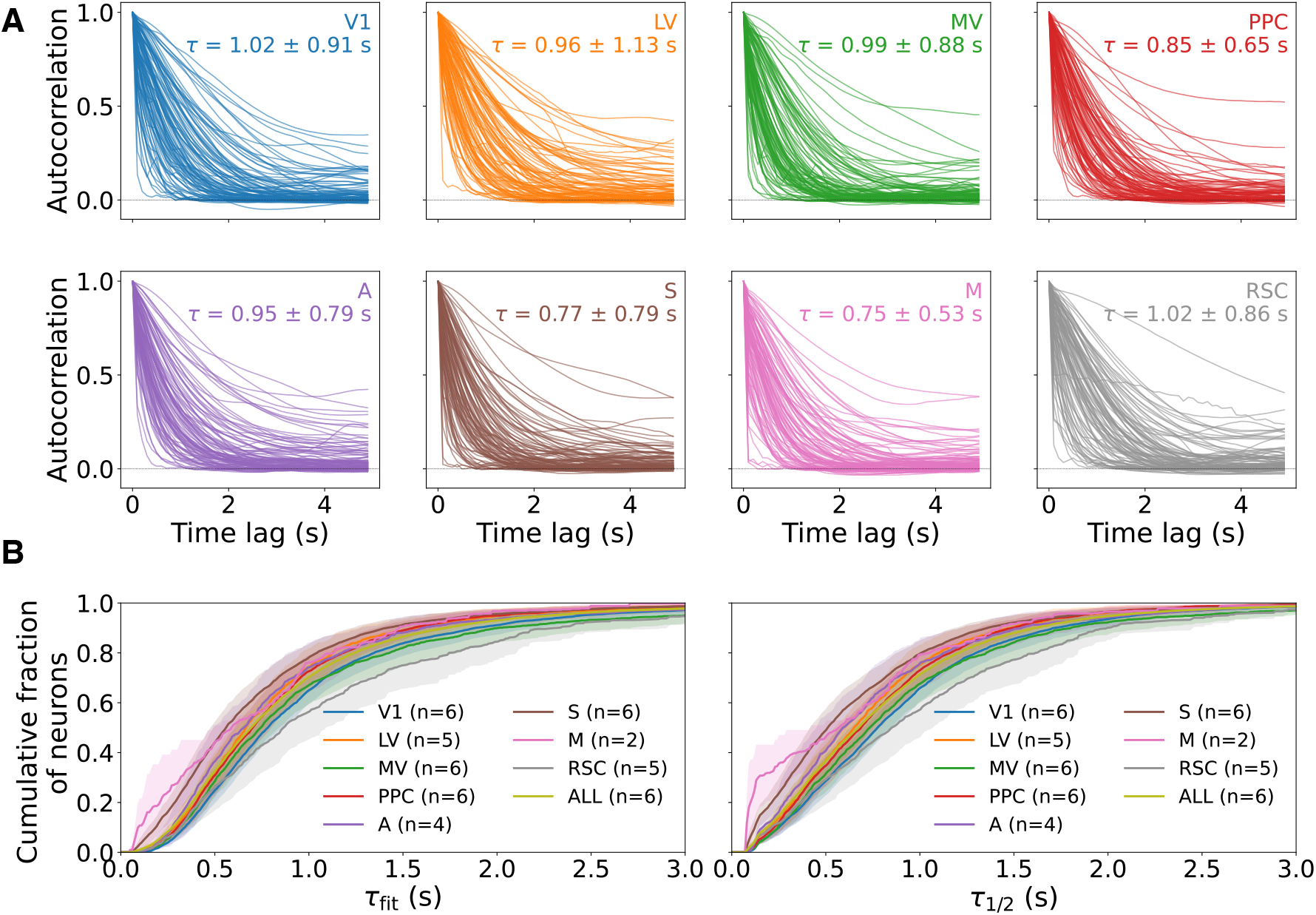
Single-neuron autocorrelation timescales across neocortical areas. **A** Example temporal autocorrelations of individual neurons in eight cortical areas: primary visual cortex (V1), lateral visual areas (LV), medial visual areas (MV), posterior parietal cortex (PPC), auditory cortex (A), somatosensory cortex (S), motor cortex (M), and retrosplenial cortex (RSC). Each panel shows (up to) 100 randomly selected neurons pooled across animals. Insets report the per-area mean *±* standard deviation of the exponential autocorrelation time constant *τ*_fit_, estimated separately for each neuron by a weighted linear fit of the log-autocorrelation function between lags 0 to 1 s. **B** Cumulative fraction of neurons whose autocorrelation time constant is at most *t*, shown for two complementary timescale estimates: the exponential-fit time constant *τ*_fit_ (left) and the half-crossing-derived time constant *τ*_1*/*2_ = *t*_0.5_*/* log 2, where *t*_0.5_ denotes the lag at which the autocorrelation crosses 0.5 (right). Solid lines show the across-animal mean and shaded bands show the standard error of the mean. An animal is included for a given area only if at least 50 of its cells in that area yield a finite time constant. Colors denote cortical areas as in panel **A**, with ALL denoting all recorded neurons pooled across areas.

**Figure S4.**
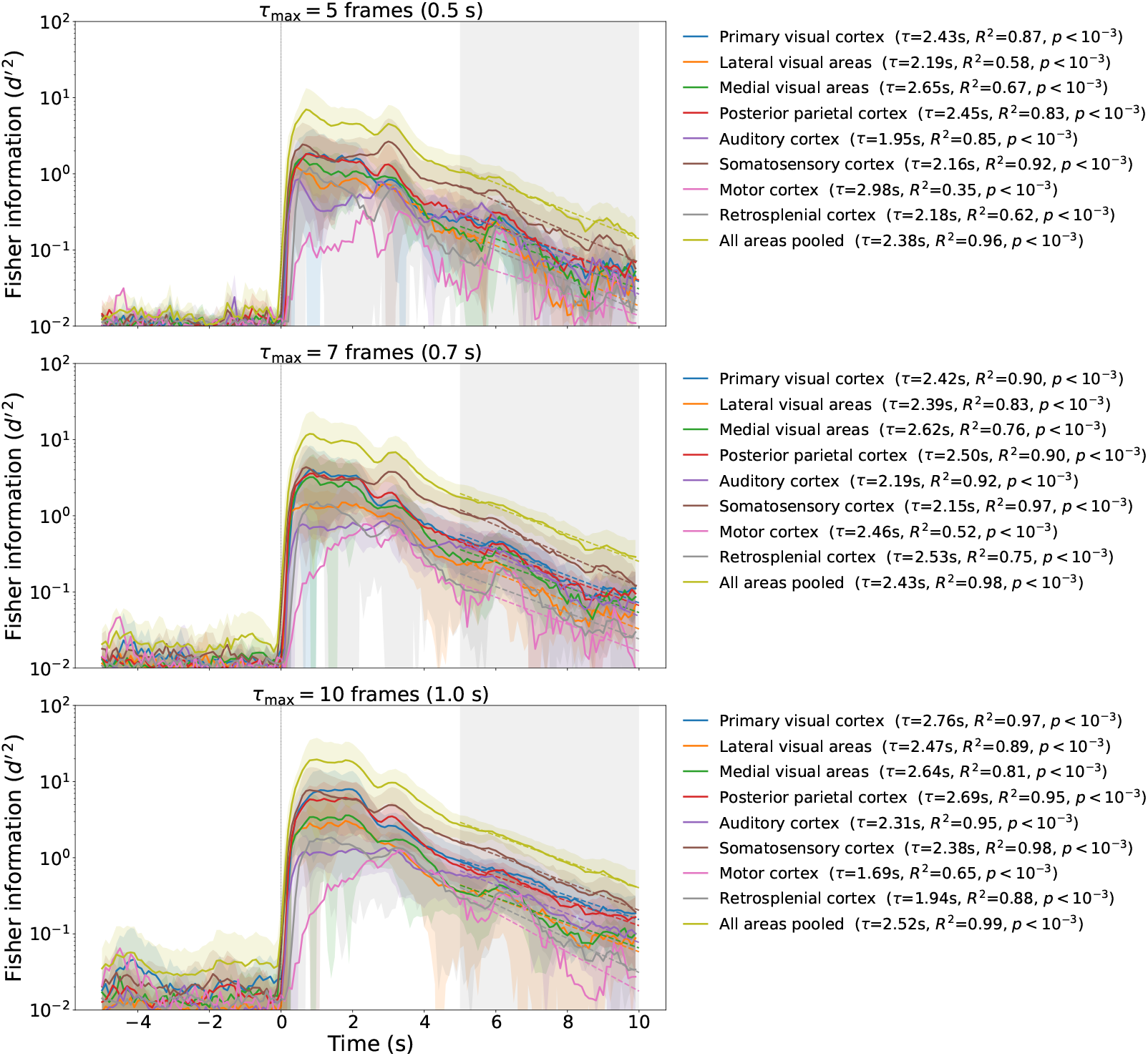
Slow decay of population Fisher information persists after excluding neurons with long single-cell autocorrelations. This figure repeats the population-level decoding analysis of Fig. 2 after restricting the analysis to cells with fast single-neuron autocorrelations. Each row corresponds to a different upper bound *τ*_max_ on the per-neuron autocorrelation timescale: *τ*_max_ = 0.5 s, 0.7 s, and 1.0 s (top to bottom). For each animal and each cortical area, a neuron is included only if both its exponential-fit autocorrelation time constant *τ*_fit_ and its half-crossing-derived time constant *τ*_1*/*2_ lie at or below *τ*_max_. After this cell selection, a PLS-LDA decoder with 15 PLS components is trained at each timepoint to discriminate trial type from population activity. Solid lines show the across-animal mean of *d*^*′* 2^ averaged over 100 train/test splits per timepoint, and shaded bands show the across-animal standard deviation. An (animal, area) pair is included only when at least 50 cells survive the timescale filter. Dashed lines show single-exponential fits to the area-mean curves over the gray-shaded window from 5 s to 10 s; the fitted decay time constant for each area appears in the legend. Across all three values of *τ*_max_, the population *d*^*′* 2^ decays with time constants comparable to those obtained without the timescale filter as in Fig. 2, indicating that the long population-level decay is not inherited from a small subpopulation of slowly varying individual neurons.

**Figure S5.**
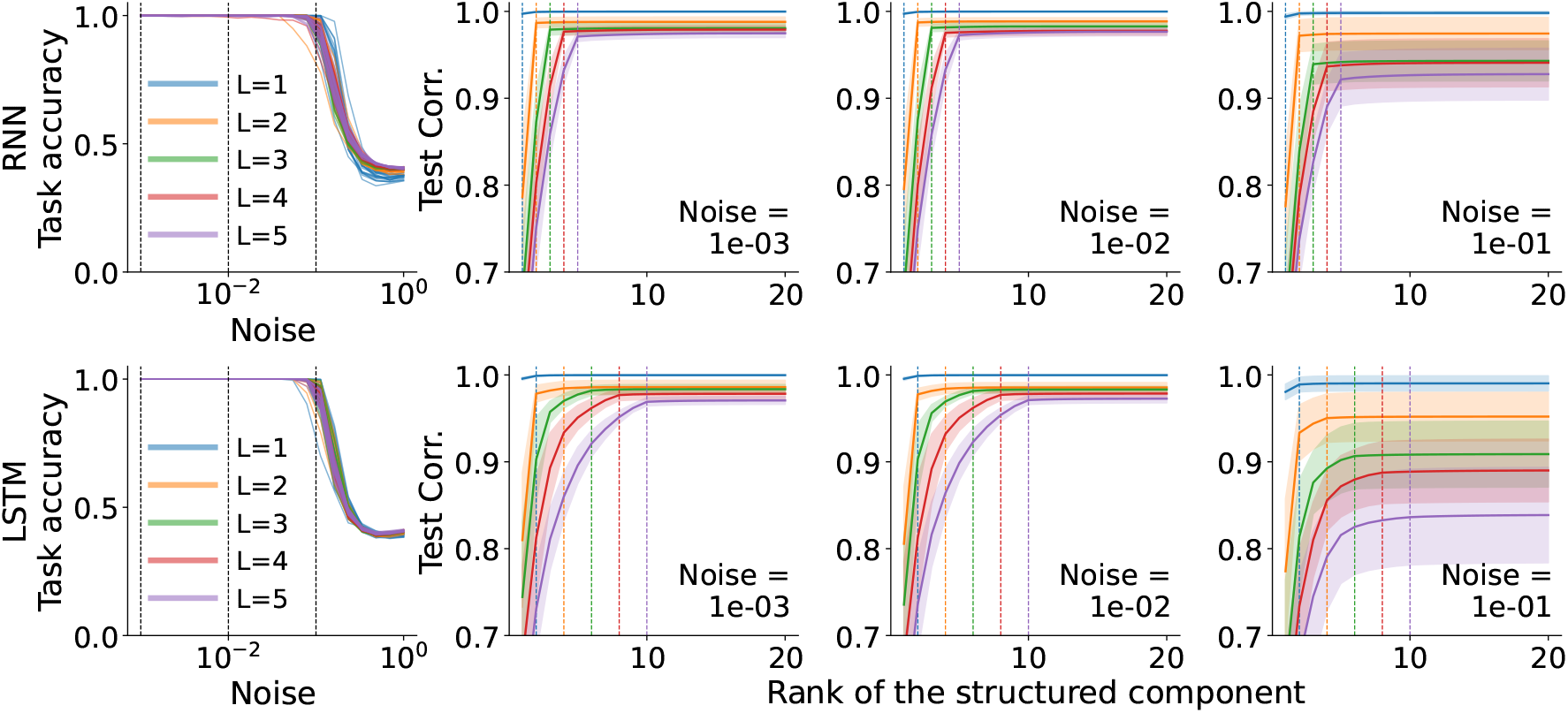
Estimation of the noise covariance matrix in task-trained RNNs. The first column shows task accuracy as a function of the magnitude of injected noise for RNNs (top) and LSTMs (bottom), for networks trained to perform *L*-bit flip-flop tasks with *L* = 1, 2, 3, 4, 5 and no noise injection during training. Vertical dashed lines mark the three noise levels used in subsequent panels. Each curve corresponds to an individual seed and colors indicate *L*. In the remaining columns, we estimate the noise covariance. We collect hidden-state activity along long flip-flop sequences, mean-center within each flip-flop state, and perform 80/20 splits to form held-in and held-out empirical covariances. We fit a low-rank-plus-diagonal model to the held-in covariance by alternating minimization. The remaining columns show the Pearson correlation between the off-diagonal entries of the fit and those of the held-out covariance, as a function of the rank of the structured component, at noise levels *σ* ∈ {10^−3^, 10^−2^, 10^−1^}. For each network, we select the best regularization strength and average over 10 independent sequences. Solid lines show the mean; shaded bands indicate *±*1 s.d. over 20 trained networks. Colored dashed vertical lines mark *L* for RNNs (top) and 2*L* for the LSTMs (bottom).

**Figure S6.**
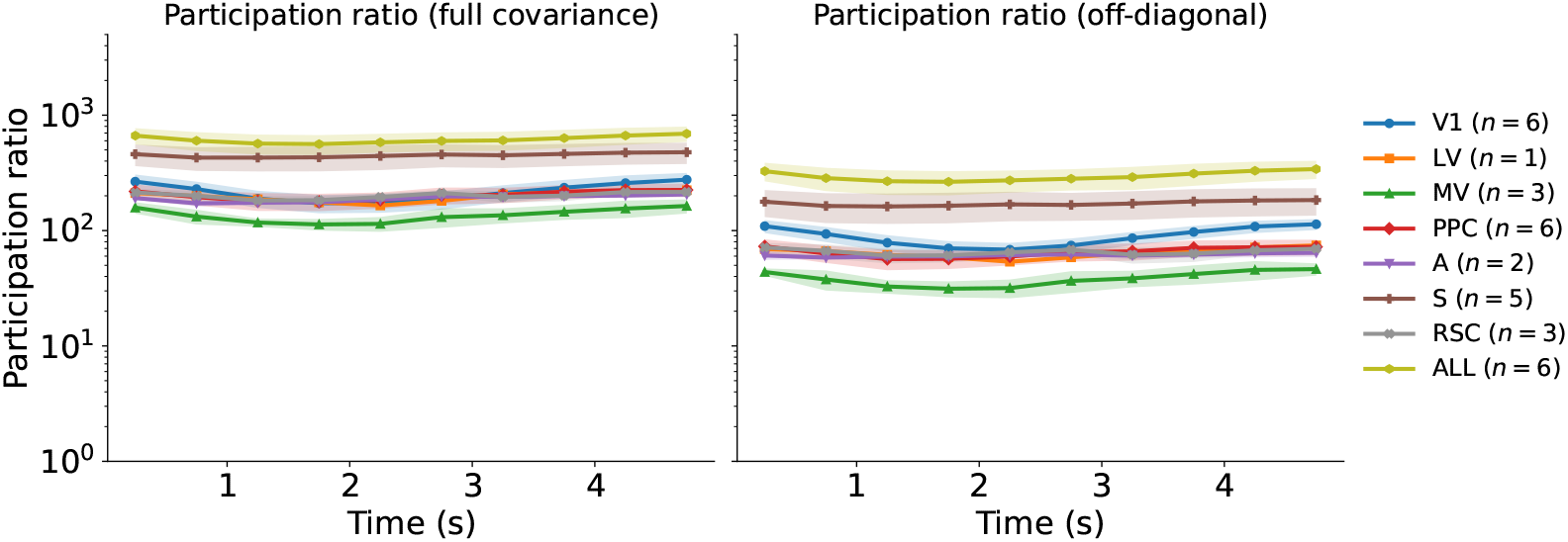
Cortical noise dimensionality across areas measured by participation ratio. Participation ratio (PR) as a function of time within a trial, computed separately for each cortical area (color-coded as in Fig. 2). *Left:* PR computed on the full noise covariance matrix. *Right:* PR computed on the off-diagonal noise covariance matrix (diagonal set to zero, using only positive eigenvalues), isolating shared variability. Traces show the mean across animals (up to *n* = 6, see legends), shaded regions denote *±*1 SEM.

**Figure S7.**
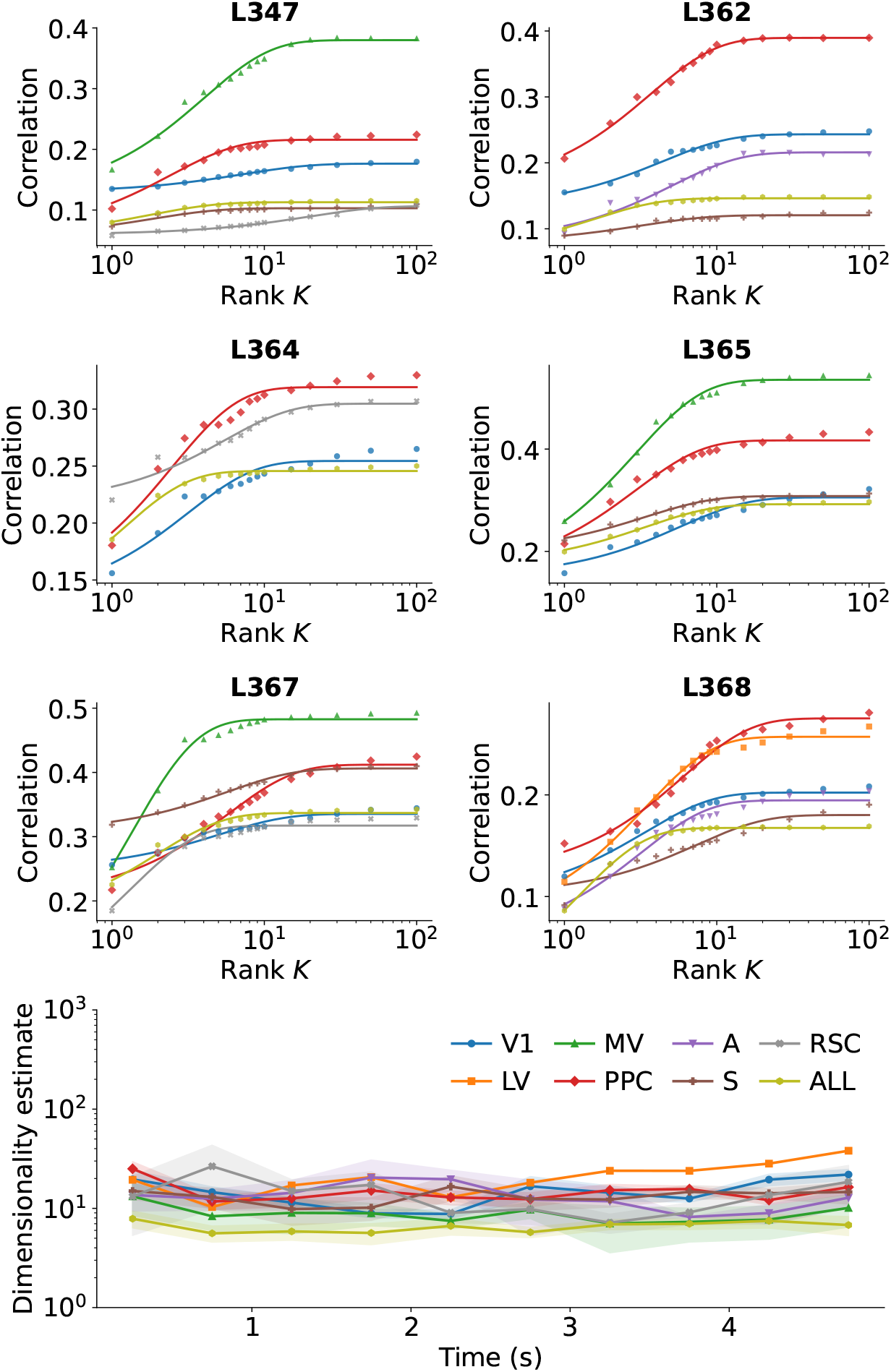
Dimensionality estimates in neocortical recordings via covariance matching. *(Top)* Exponential fits to the test-set correlation as a function of rank *K* for each animal and brain region (colors) at a representative time window during the task (0.5-1 s after stimulus onset). For each of 20 random 80/20 train/test splits of trials, we compute the covariance matrices of the held-in and held-out populations, fit a diagonal-plus-rank-*K* model to the training covariance, and evaluate its correlation with the test covariance; the value is then maximized over 10 regularization strengths sampled logarithmically distributed from *λ* ∈ [10^−2^, 10^2^] and averaged across splits (data points). Curves show the fitted saturating exponential *A*(1 − *e*^−*K/τ*^) + *c*, see Appendix S4.3 for details. *(Bottom)* The plot shows the dimensionality estimates, *d*_CM_ = 3*τ*, as a function of time across the trial, averaged across animals (mean *±* SEM).

**Figure S8.**
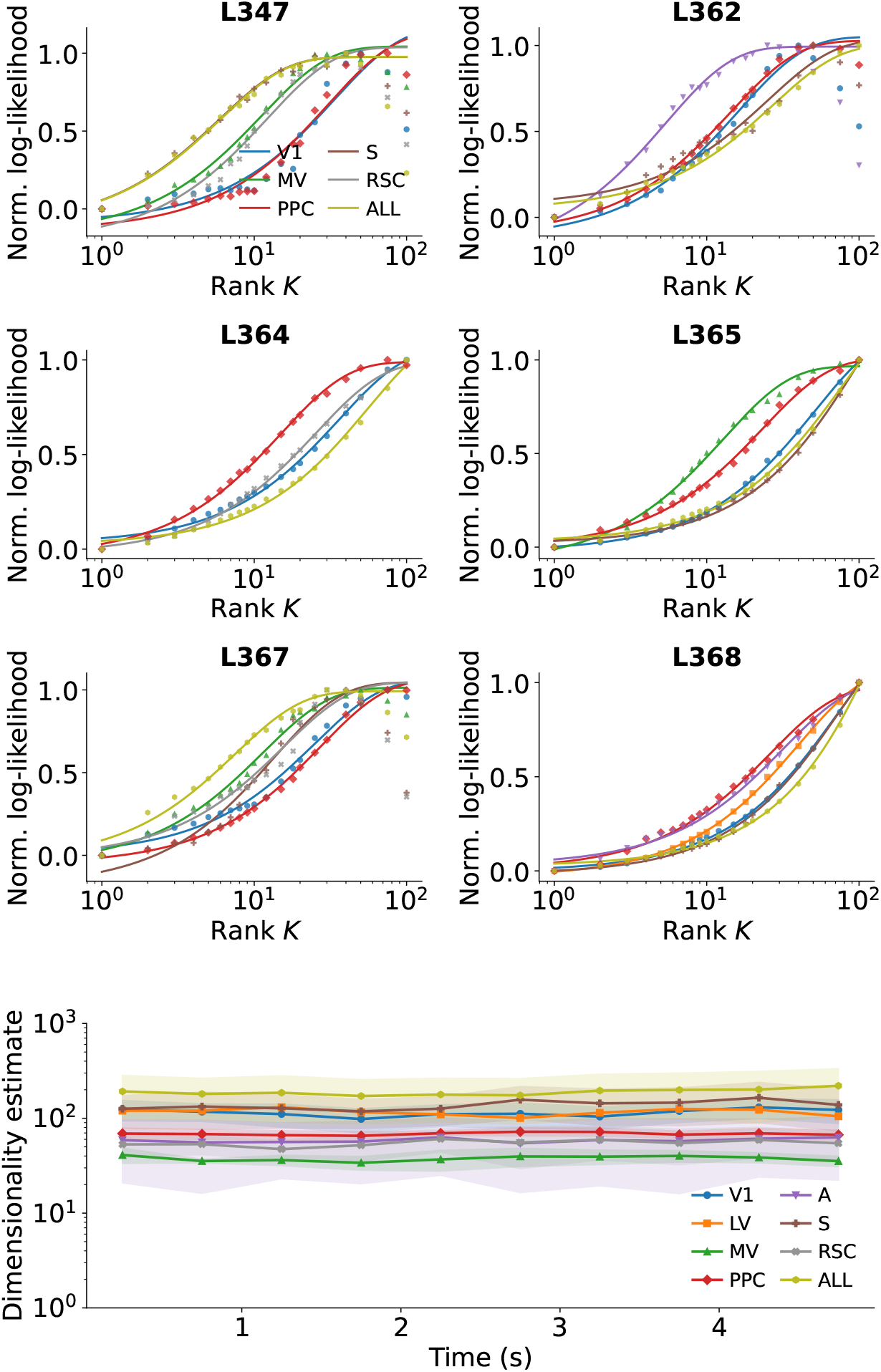
Dimensionality estimates in neocortical recordings via factor analysis. *(Top)* Exponential fits to the test-set log-likelihood as a function of the number of latent factors *K* for each animal and brain region (colors) at a representative time window during the task (0.5-1 s after stimulus onset). For each of 20 random 80/20 train/test splits of trials, we fit a factor analysis model with *K* components to the held-in data and evaluate the median log-likelihood on held-out samples; values are then averaged across splits and normalized per fitting instance (subtract minimum, divide by absolute maximum) to place curves on a common scale (see data points). Curves show the fitted saturating exponential *A*(1 − *e*^−*K/τ*^) + *c*, see Appendix S4.3 for details. *(Bottom)* The plot shows the dimensionality estimates, *d*_FA_ = 3*τ*, as a function of time across the trial, averaged across animals (mean *±* SEM).

**Figure S9.**
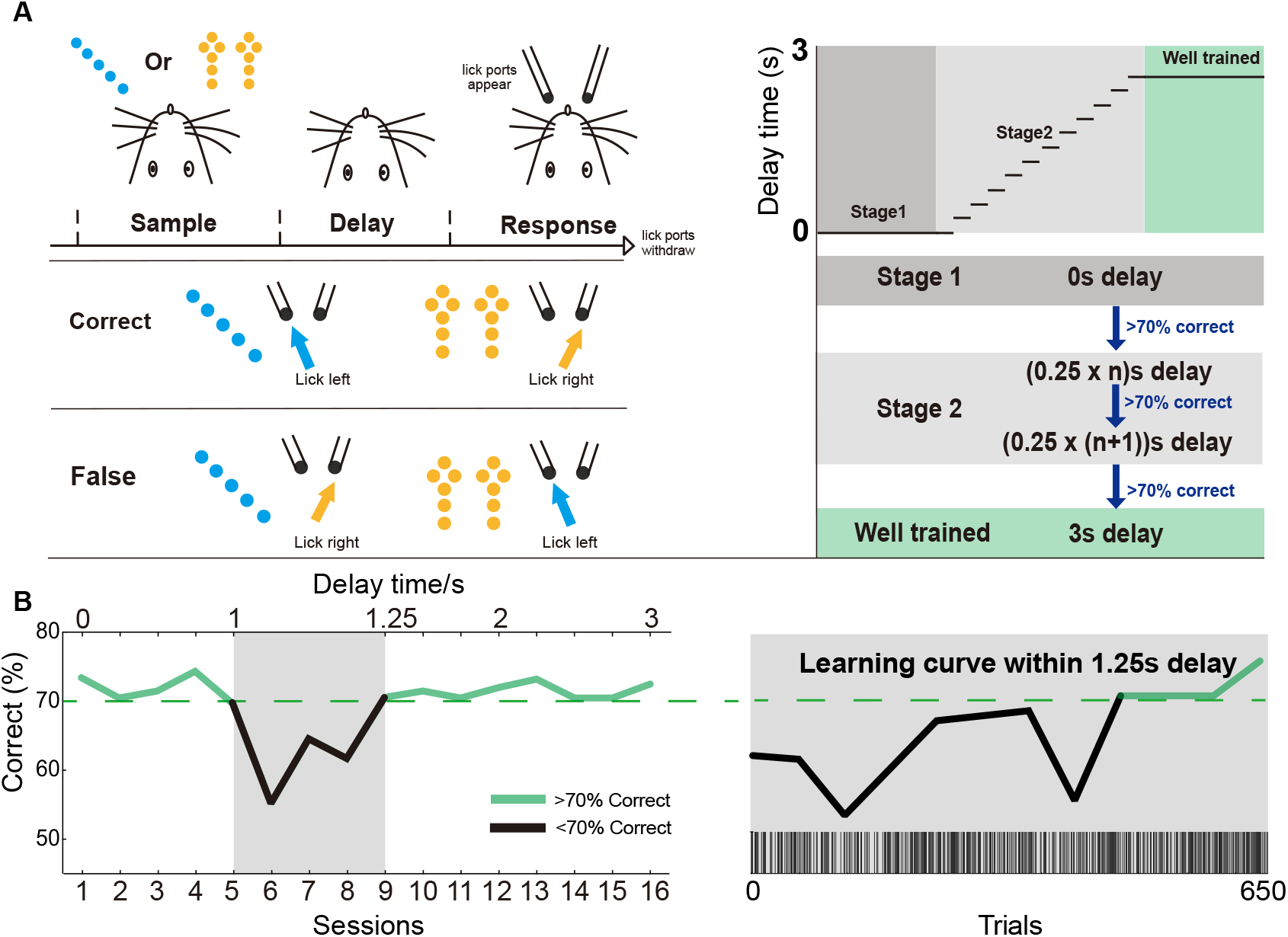
The training paradigm for testing the short-term memory limits in mice. **A** Schematics of the training procedure. Mice were presented with a blue or yellow visual cue, followed by a delay period and a response window during which the lick ports became available. Animals were first trained with 0s delay (Stage 1), and when reached 70% correct, advanced to Stage 2, in which delay duration was increased in 0.25s increments once performance exceeded 70% correct, until stable performance was achieved at 3s delay. **B** Representative behavioral performance across training sessions from a single mouse. Left, session-by-session accuracy during progressive delay training. Green segments indicate sessions with performance above 70%, whereas black segments indicate sessions below 70%. The shaded region denotes the 1-1.25s delay range. Right, learning curve at the 1.25s delay plotted as a function of trial number. Black bars at the bottom indicate correct trials. The solid line corresponds to the correct rate for each consecutive block of 50 trials.

**Figure S10.**
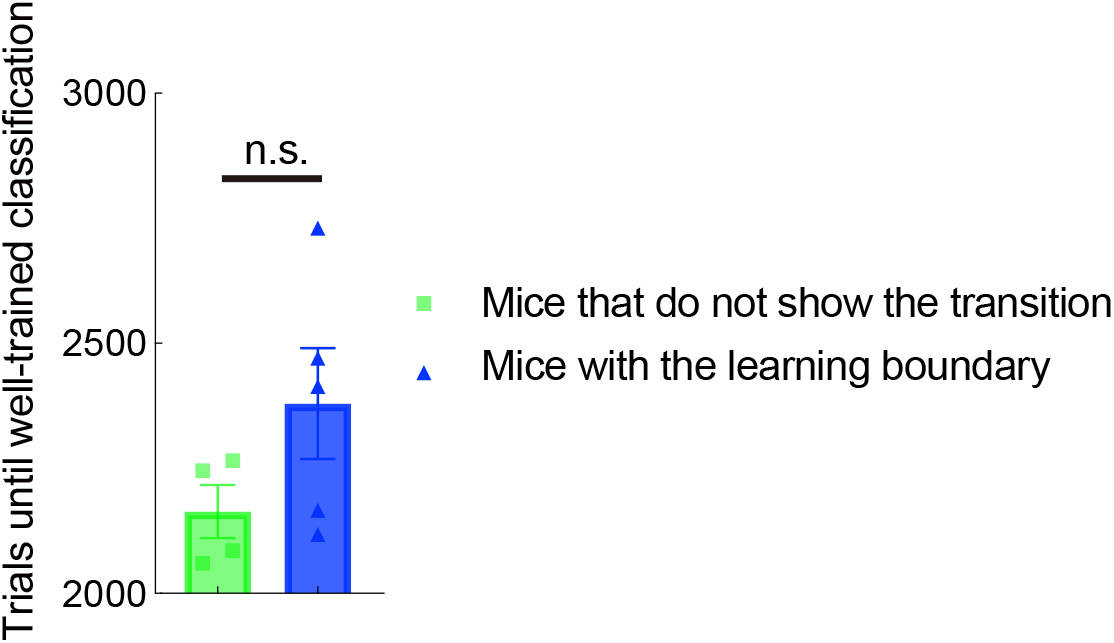
Number of trials before the incremental delay training. The bar plots show the number of trials needed for the mice to be considered well-trained before the increasing delays start. Each dot corresponds to a mouse from Fig. 4**B-C**, divided into two groups: those that did not show a clear learning boundary vs those that did. Although the former required fewer trials to reach criterion, the group difference did not reach statistical significance (*p* = .152, unpaired t-test).

**Figure S11.**
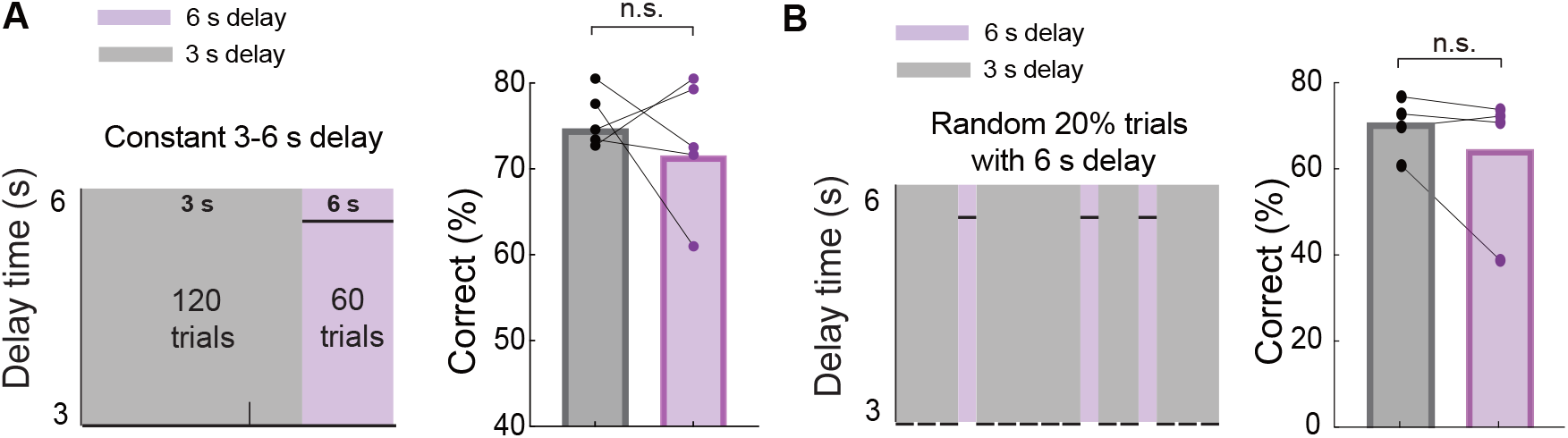
Working memory performance generalizes to novel 6s delays without additional training. **A** Behavioral performance in a session with 120 trials with 3s delay, followed by 60 trials with 6s delay. Mice encountered 6s delay for the first time. Dots represent individual mice; lines connect paired observations from the same animal. No significant difference was detected between the two delay conditions (paired test, *p* = 0.563, n=5). **B** Behavioral performance in a session with 20% randomly assigned trials with 6s delay. Mice encountered 6s delay for the first time. Dots represent individual mice; lines connect paired observations from the same animal. Performance in 6s delay trials showed a trend towards reduction compared to 3s delay, which did not reach significance (paired test, *p* = 0.313, n=4).

## Footnotes

1 These random variables capture both neuron-wise processes and the network connections. For instance, the sample 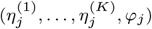 explains both the neuron-wise nonlinearity and the synaptic parameters associated with the neuron *j*.

